# Coordinated shifts in gene expression and regulation during mole-rat evolution

**DOI:** 10.1101/2025.11.24.690111

**Authors:** Maëlle Daunesse, Eulalie Liorzou, Elise Parey, Diego Villar, Camille Berthelot

## Abstract

Changes in gene expression and regulation are central to mammalian phenotypic evolution. Yet, distinguishing adaptive gene expression shifts from neutral divergence remains challenging. Here, we integrate comparative transcriptomics and regulatory genomics to investigate how natural selection has shaped gene expression in African mole-rats, a group of subterranean rodents with phenotypic adaptations to underground environments. Using RNA-seq from liver and heart in two mole-rat species (naked mole-rat and Damaraland mole-rat) and two rodent outgroups (mouse and guinea pig), we leveraged phylogenetic models of expression evolution and identified hundreds of genes whose transcriptional levels have experienced accelerated evolution in each tissue and mole-rat species. These lineage-specific shifts account for only a small fraction of gene expression differences identified by classical differential expression analysis between species, underscoring the importance of phylogeny-aware inference to disentangle accelerated evolution from drift. To connect expression divergence with regulatory evolution, we integrated transcriptomic profiles with cis-regulatory landscapes. Genes with lineage-specific expression shifts displayed concordant changes in cis-regulatory activity, particularly at promoters, and the magnitude of expression divergence increased with the number of shifted cis-regulatory elements. Our results demonstrate that adaptive shifts in mole-rat gene expression are mirrored by regulatory evolution, providing genome-wide coordinated evidence of accelerated evolution on expression and regulation during mammalian evolution. Our approach thereby prioritises candidate loci that may have shaped adaptations specific to mole-rat physiology, including metabolic rewiring and stress responses.

## Introduction

Changes in gene expression are a major substrate of phenotypic evolution. In mammals, expression levels are generally more conserved than regulatory activity, reflecting the action of stabilising selection to maintain cellular transcriptomic profiles (Brawand et al. 2011; Necsulea and Kaessmann 2014; Breschi et al. 2016; Berthelot et al. 2018; Cardoso-Moreira et al. 2019). By contrast, cis-regulatory elements (CREs), in particular enhancers, evolve rapidly with frequent gains and losses between lineages (Cotney et al. 2013; Reilly et al. 2015; Villar et al. 2015; Roller et al. 2021; Phan et al. 2025). This discrepancy raises a central question in evolutionary genomics: to what extent do regulatory changes translate into functional divergence in gene expression, and when do both expression and regulation layers carry concordant molecular signatures of selection?

Addressing this question is challenging for multiple reasons. First, expression and regulatory evolution have been frequently analysed independently for practical reasons, and few studies have collected matched transcriptomic and epigenomic data across multiple species (Berthelot et al. 2018; Sarropoulos et al. 2025; Foissac et al. 2019; Reilly et al. 2015; Halstead et al. 2020) Second, models to detect accelerated evolution acting on gene expression and regulation activity remain an active area of research, in particular to distinguish between evidence of selection and relaxation (Price et al. 2022; Rohlfs and Nielsen 2015; Zambon et al. 2025). Third, connecting divergence in gene expression and regulation with functional differences between species remains difficult (Romero et al. 2012), and ideally requires a model system where the molecular basis of phenotypic adaptations is well characterised. As a result, how regulatory changes articulate to drive adaptive evolution of gene expression remains poorly understood.

Phylogenetic comparative methods provide a framework to detect putative selection on quantitative traits such as gene expression. In particular, Brownian motion and Ornstein–Uhlenbeck models have long been used to distinguish neutral drift from stabilising selection across a variety of continuous traits, including molecular phenotypes such as gene expression (Felsenstein 1985; Hansen 1997; Khaitovich et al. 2006; Wong et al. 2015; Price et al. 2022).

The Expression Variance and Evolution (EVE) model extended this framework by incorporating inter-and intra-species variance, enabling branch-specific tests for shifts in optimal expression levels(Rohlfs and Nielsen 2015). Applications of such models have revealed lineage-specific shifts in mammalian transcriptomes (Chen et al. 2019; Brawand et al. 2011; Breschi et al. 2016), but integration with regulatory evolution is still limited.

Here, we apply a comparative phylogenetic framework to African mole-rats and two rodent outgroups to investigate genome-wide shifts in gene expression and their corresponding regulatory activity. By integrating transcriptomic and regulatory data, we seek to distinguish putative adaptive shifts from divergence consistent with neutral drift, and to highlight candidate loci where both layers of molecular evolution support the same evolutionary trajectory.

Mole-rats are a diverse group of over 20 subterranean rodent species found across sub-Saharan Africa. They exhibit remarkable adaptations to underground life and a broad range of social behaviours, from solitary to eusocial (Cox et al. 2020). The naked mole-rat (*Heterocephalus glaber*) and the Damaraland mole-rat (*Fukomys damarensis*) are both eusocial and long-lived, reaching lifespans of 37 and 26 years despite small body sizes (Dammann et al., n.d.; Can et al. 2022; Begall et al. 2021). Among their reported adaptations, these species show high tolerance to hypoxia (Ivy et al. 2020), hypercapnia (Park and Reznick 2023), and oxidative stress (Lewis et al. 2013), with the naked mole-rat also displaying cancer resistance (Buffenstein et al. 2022). While both exhibit molecular traits linked to subterranean adaptation, the naked mole-rat shows more pronounced evolutionary divergence (Faulkes and Bennett 2021). In sum, phenotypic adaptation in African mole-rats has been well described at the morphological and physiological levels (Faulkes and Bennett 2021; Buffenstein et al. 2022) making them a good model system for our study.

Our results show that genes with expression shifts identified by phylogenetic modelling are significantly enriched for concordant changes in their regulatory landscapes, particularly at promoters, and that the magnitude of expression divergence scales with the number of associated regulatory shifts. These genes implicate pathways related to metabolism, stress responses, and cardiac physiology, align with phenotypic traits previously described in mole-rats, and indicate potentially adaptative roles for the candidate loci we identify. Together, this work provides genome-wide evidence that regulatory and expression evolution exhibit concordant evolutionary shifts consistent with signatures of selection and establishes a phylogeny-aware framework to prioritise potentially adaptive loci during mammalian evolution.

## Results

### Comparative transcriptomics of liver and heart gene expression in mole-rats

To investigate gene expression changes during mole-rat evolution, we generated bulk RNA sequencing libraries from two tissues with distinct metabolic functions, liver and heart, in two mole-rat species: naked mole-rat (*Heterocephalus glaber*; Hgla) and Damaraland mole-rat (*Fukomys damarensis*; Fdam), and two outgroups: guinea pig (*Cavia porcellus*; Cpor), and mouse (*Mus musculus*; Mmus) (**Figure 1A**). We generated libraries from 28 distinct samples, and complemented this dataset with publicly available RNA-seq from previous studies (Berthelot et al. 2018; Bens et al. 2018; Faulkes et al. 2024). The collated data includes 5 to 8 biological replicates per species and tissue, resulting in a robust dataset for evolutionary modelling.

**Figure 1:**
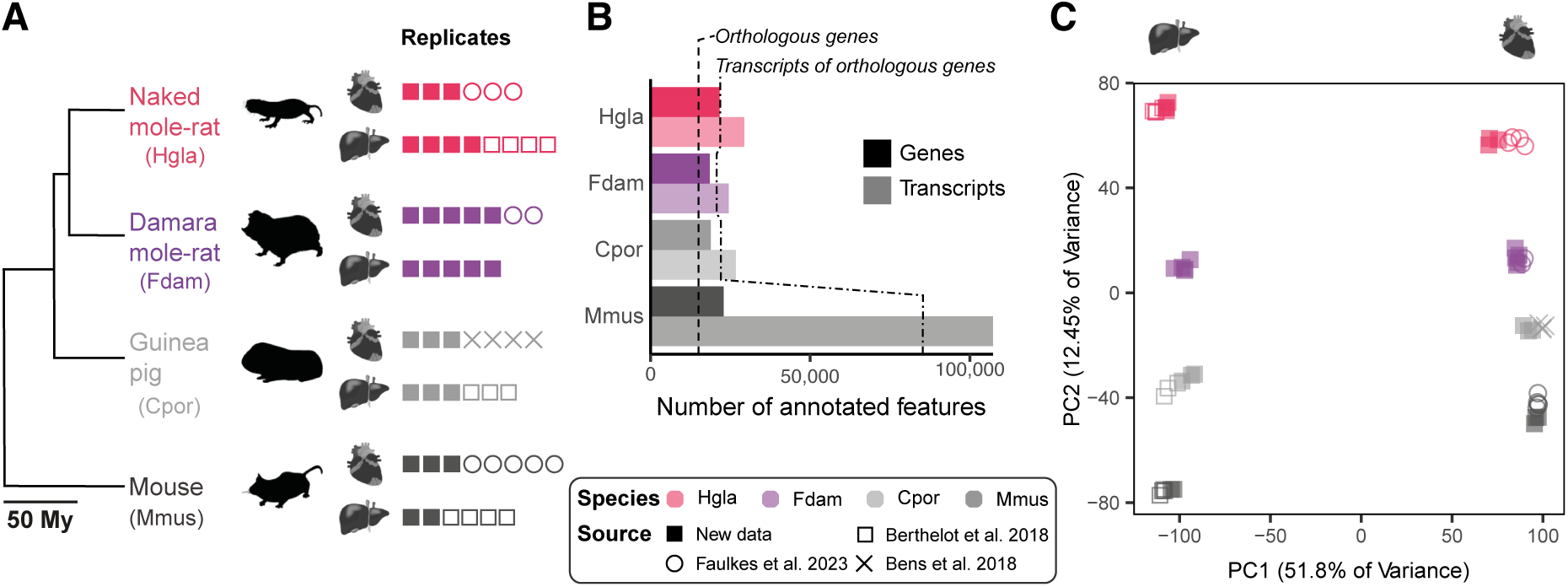
Experimental design overview. **(A)** This study includes RNA-seq libraries from two tissues (liver and heart) and four species: naked mole-rat (Hgla), Damaraland mole-rat (Fdam), guinea pig (Cpor), and mouse (Mmus), with 5–8 biological replicates per species and tissue. **(B)** Non-model species (Hgla, Fdam, and Cpor) have similar numbers of genes and transcripts, while the mouse (Mmus) has a higher quality annotation, particularly in terms of annotated transcripts. A total of 14,408 orthologous genes were identified across the four species, with respectively 20,905, 19,685, 21,102 and 81,378 annotated transcripts in Hgla, Fdam, Cpor and Mmus. **(C)** Principal component analysis of variance-stabilised, normalised gene expression levels shows strong clustering by tissue along PC1 and by species along PC2, indicating high reproducibility and minimal batch effects.

To compare gene expression levels across the four species, we identified 14,408 one-to-one orthologous genes by combining orthology annotations from Ensembl v102 (Yates et al. 2020) and from the Zoonomia consortium (Christmas et al. 2023) (**Methods**, **Figure S1**; **Table S1**). These one-to-one orthologous genes include a most of the annotated protein-coding genes in all four species (Hgla: 70%; Fdam: 81%, Cpor: 80% and Mmus: 66%). In each species, we used pseudo-mapping with Salmon (Patro et al. 2017) to map RNA-seq reads to annotated transcripts (**Methods**). Differences in transcript diversity and annotation quality can introduce systematic biases: indeed, our study species vary significantly in number of annotated transcript isoforms (**Figure 1B**), with mouse having the most comprehensive annotation (**Figure S2**). To correct for these sources of variation, we normalised transcript counts using the effective transcript length estimated by Salmon during quantification (Patro et al. 2017). This metric adjusts for transcript length and accounts for sequence-specific and positional bias, thereby correcting for differences in transcript diversity and annotation quality across species (**Methods**). Our results show that this normalisation strategy significantly improves replicate consistency and integration compared to normalising by the theoretical gene length (**Figure S3**). We then aggregated transcript-level estimates into gene-level quantifications and normalised for library size (**Table S2**).

To assess global expression patterns and dataset consistency, we performed a principal component analysis (PCA) on the orthologous gene expression matrix after variance-stabilising transformation (**Figure 1C**). Samples separated primarily by tissue along PC1 (51.8% of variance), and by species along PC2 (12.45%), indicating strong biological signal and minimal batch effects. Replicates clustered tightly, including across data sources, reflecting high reproducibility of gene expression levels within species and tissues. Additionally, the distribution of species along PC2 is consistent with their phylogenetic relationships (**Figure 1C**).

### Phylogenetic modelling of lineage-specific gene expression shifts in mole-rats

To assess accelerated evolution of gene expression levels in mole-rats, we next sought to identify genes exhibiting lineage-specific expression changes in either the ancestral mole-rat branch (Ancestral), or one of the terminal branches leading to the naked mole-rat (Hgla) and the Damaraland mole-rat (Fdam). We first performed differential expression (DE) analysis contrasting gene expression levels between the foreground and background species in liver and heart (**Figure 2A**; **Table S3**), and identified thousands of differentially expressed genes per contrast, ranging from 4,721 to 7,960. However, differential expression does not account for the shared evolutionary history of our study species and cannot discriminate between accelerated evolution on a specific branch and divergence consistent with neutral drift (**Figure 2B**). To refine this list, we applied the Expression Variance and Evolution (EVE) model (Rohlfs and Nielsen 2015), a phylogenetic comparative method that models expression as a quantitative trait evolving under an Ornstein-Uhlenbeck process (**Figure 2B**; **Table S3**). EVE explicitly accounts for phylogenetic relationships and implements a likelihood ratio test to distinguish between both evolutionary scenarios.

**Figure 2:**
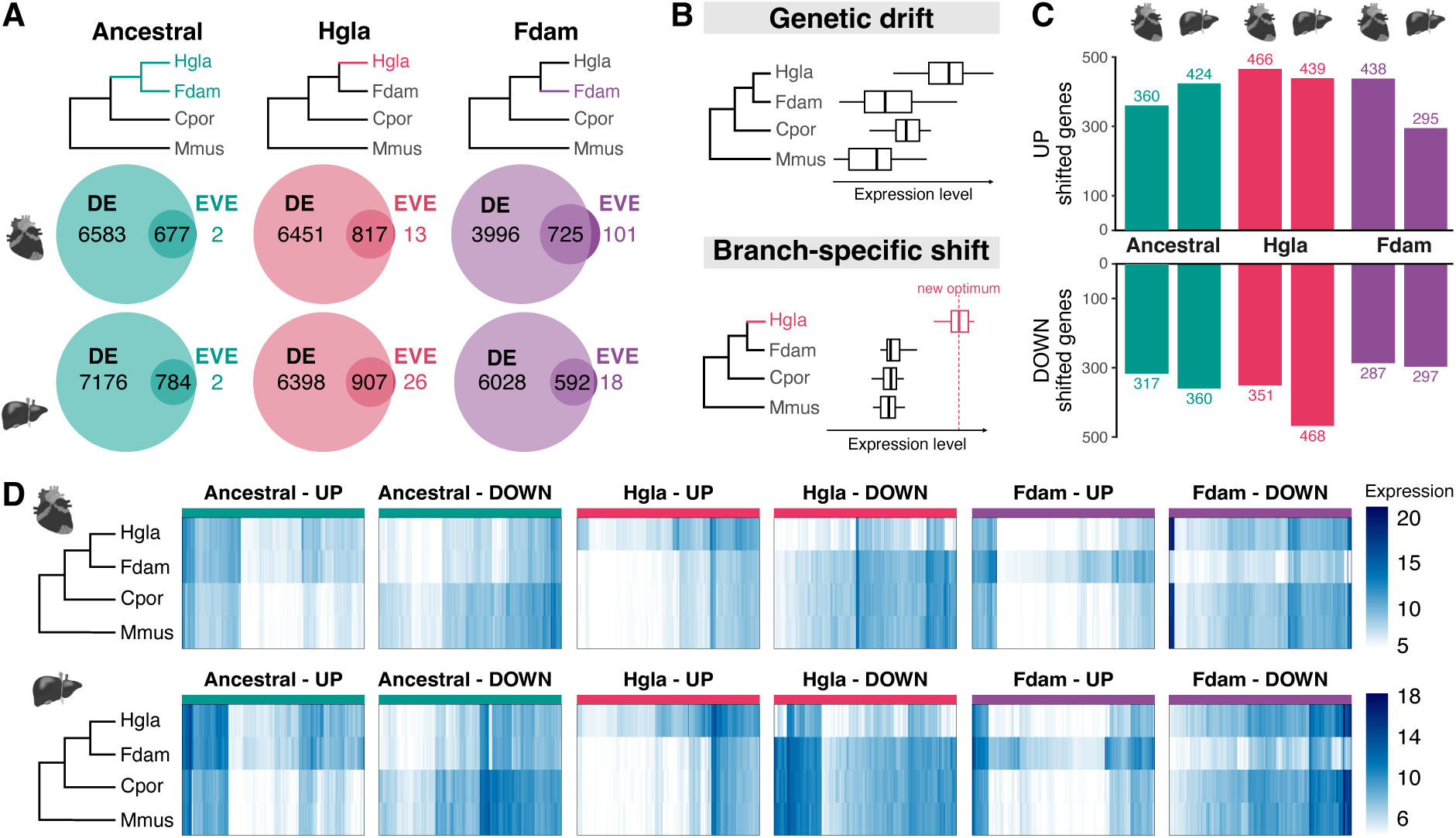
Phylogenetic modelling of lineage-specific gene expression shifts in mole-rats. **(A)** Numbers of differentially expressed (DE) genes (left circles) and lineage-shifted genes identified by the EVE model (right circles) for the ancestral mole-rat branch, the naked mole-rat (Hgla), and the Damaraland mole-rat (Fdam). Overlap between DE and EVE sets (intersection) defines the final set of shifted genes. **(B)** Conceptual framework: differential expression can result from either genetic drift (top) or branch-specific shifts consistent with adaptive evolution (bottom). EVE (Rohlfs and Nielsen 2015) distinguishes between these scenarios by modelling expression evolution under an Ornstein–Uhlenbeck process. **(C)** Numbers of up-and down-regulated genes per branch and tissue after intersecting DE and EVE results. **(D)** Heatmaps showing variance-stabilised, normalised expression values for shifted genes in each branch and tissue.

The intersection of differentially expressed genes (DE) and lineage-shifted genes (EVE) revealed that most differentially expressed genes are in fact consistent with neutral divergence, and only a subset (9-15%) display evidence of accelerated evolution (**Figure 2A**; **Table S3**). Conversely, almost all genes that experienced an episode of accelerated evolution as detected by EVE show significant differences in expression between background and foreground species. For further analysis, we restrained the gene set to genes supported by both methods, resulting in 295 to 466 up-regulated and 287 to 468 down-regulated genes per branch and tissue (**Figure 2C**). For each branch, species within the tested branch displayed consistent expression changes, whereas background species showed conserved expression levels (**Figure 2D**). This pattern reinforces the interpretation that the detected shifts reflect lineage-specific evolutionary dynamics consistent with accelerated evolution rather than broader transcriptional noise.

### Lineage-specific shifts in mole-rat gene expression are tissue-specific and associate to previously proposed mole-rat adaptations

We then investigated whether shifted genes overlap between branches and tissues. We found that the signal was highly branch-specific: no gene was identified as shifted in more than one branch within a given tissue. This likely reflects the relatively small number of species in our phylogenetic sampling, which does not allow to detect multiple changes in the tree. However, about 20% of all genes shifted in liver were also detected as shifted in heart for all tested branches, which represents a much higher overlap than expected by chance (hypergeometric test with BH adjustment: Ancestral, p = 2.8 × 10⁻⁵³; Hgla, p = 3.2 × 10⁻⁸⁶; Fdam, p = 6.5 × 10⁻⁷⁹; **Figure 3A**). Further, when a gene was shifted in both tissues, the direction of change (up-or down-regulated) was consistent in most cases (**Figure 3B** and **S4**) and the magnitude of shift was highly correlated (Pearson R between 0.74 and 0.96; **Figure S5**), suggesting that a proportion of gene expression changes reflect ubiquitous evolutionary pressures rather than tissue-specific adaptations.

**Figure 3:**
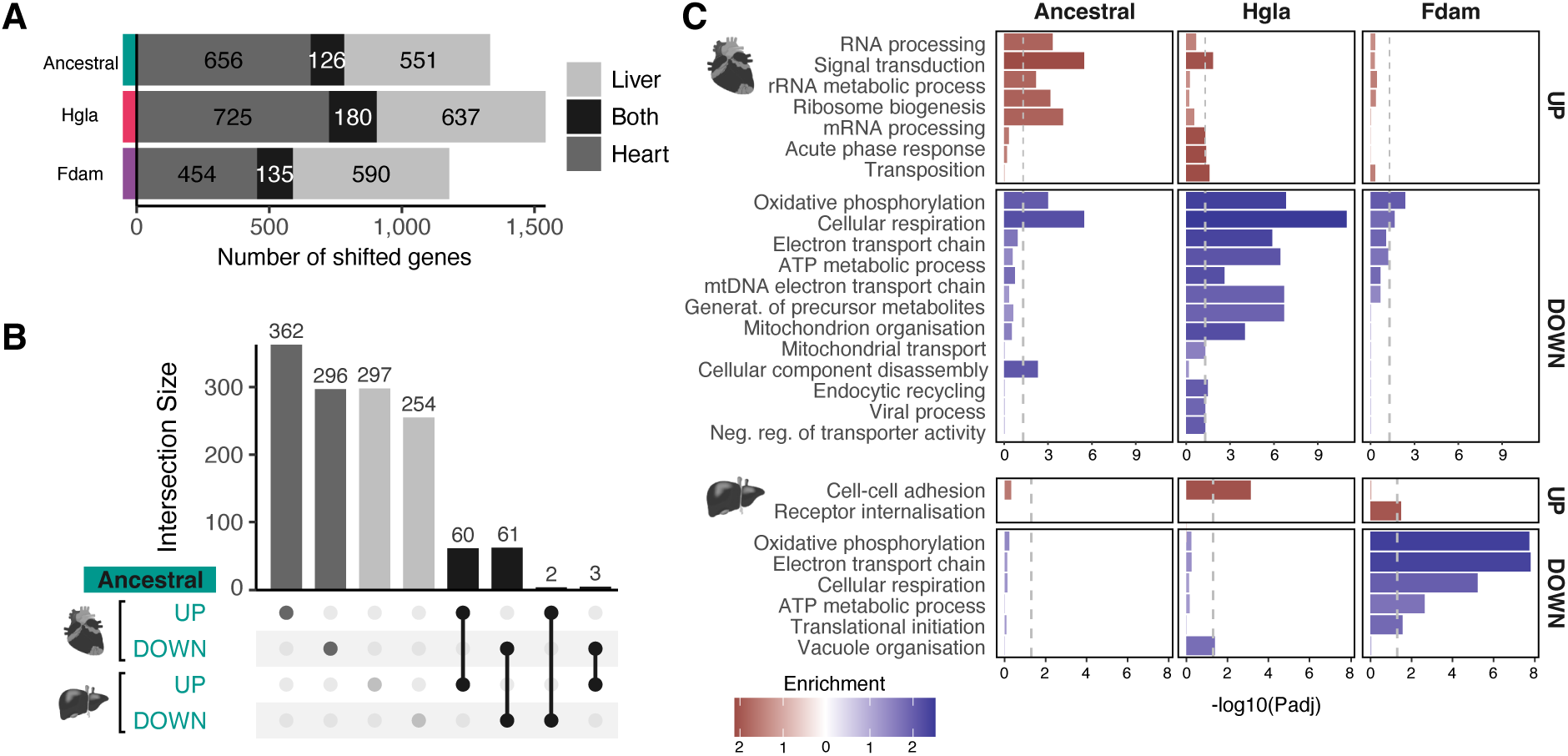
Overlap between tissues and functional enrichment of lineage-shifted genes. **(A)** Numbers of shifted genes detected in liver only (grey), heart only (dark grey), or both tissues (black) for the ancestral mole-rat branch, the naked mole-rat (Hgla), and the Damaraland mole-rat (Fdam). **(B)** Intersection analysis showing the number of genes shifted in both tissues within each branch. Bars indicate the numbers of shifted genes in the intersection; connected dots indicate the type of overlap. **(C)** Gene set enrichment analysis (GSEA) of shifted genes for each branch and tissue, using GO Biological Processes. The significantly enriched terms for up-regulated (UP) and down-regulated (DOWN) genes are shown, coloured by normalised enrichment score (red: enriched in up-regulated genes; blue: enriched in down-regulated genes).

To assess putative biological functions of lineage-specific expression shifts, we performed gene set enrichment analyses (GSEA; **Table S4**) using GO Biological Processes (**Figure 3C**), and Reactome, KEGG, and Hallmark collections (**Figure S6**). Across branches and tissues, a consistent signal emerged for the down-regulation of oxidative metabolism. In GO Biological Processes, “oxidative phosphorylation” and “cellular respiration” were significantly enriched among down-regulated genes in most branch–tissue combinations (including *Ndufa4, Uqcr10* and *Cox7c*; **Figure S7**). Concordantly, Reactome terms such as “respiratory electron transport” supported this metabolic downshift. In the Hallmark collection, a broad swathe of genes associated with “adipogenesis” was significantly down-regulated in the ancestral and naked mole-rat branch for the heart (Hallmark, including *Lipe, Lpl* and *Mdh2)*, consistent with previous reports of reduced lipid storage and lipid metabolism in mole-rats in favour of glycogen metabolism (Bundgaard et al. 2024; Faulkes et al. 2024). These gene expression shifts are consistent with a coordinated metabolic reorganisation affecting both lipid and oxidative pathways in mole-rats, as proposed previously (Heinze et al. 2018; Farhat et al. 2020; Yap et al. 2022).

Branch-and tissue-specific enrichments highlighted additional biological processes as having experienced episodic accelerated evolution in mole-rats that aligns with previously described adaptations. In the ancestral branch, gene expression shifts included up-regulation of RNA processing functions, with significant enrichment for “rRNA processing” across both tissues (including *Nop9*, *Pdcd11, Bud23* and *Exosc10*; **Figure S7**) and “translation” in liver (Reactome, including *Rpl27* and *Rps11*; **Figure S7**). Previous work has reported mole-rat alterations in rRNA fragmentation and translational fidelity (Azpurua et al. 2013; Gutierrez-Vargas et al. 2025), and our results suggest accelerated evolution of gene expression levels across RNA processing genes in mole-rats. Conversely, “cardiac muscle contraction” was significantly down-regulated in the ancestral branch for the heart (Reactome, including *Atp1b3, Fxyd2* and *Tpm4*; **Figure S7**). These lineage-specific shifts are consistent with previously reported changes in cardiac contractility (Grimes et al. 2014; 2017), and our phylogeny-aware results indicate down-regulated gene expression is favoured in this biological process.

In sum, lineage-specific expression shifts were largely restricted to either heart or liver tissues and associated with biological processes aligned with their potential roles in phenotypic adaptation.

### Changes in regulatory activity around TSSs correlate with expression shifts

We next sought to assess how gene expression shifts interplay with changes in tissue-specific regulatory landscapes across mole-rats (Parey et al. 2023). To explore how gene expression shifts across species are operated by changes in regulatory elements, we integrated cis-regulatory element (CRE) activity with our RNA-seq expression dataset, using histone modification ChIP-seq data from our previous study (Parey et al. 2023). As described in Parey et al., putative active promoters were defined as regions with both H3K4me3 and H3K27ac enrichments, and putative active enhancers as regions with H3K27ac enrichment only. We then associated each one-to-one orthologous gene with a regulatory score, calculated as the sum of normalised fold-enrichments over input of H3K27ac ChIP-seq data at CREs located within a set distance from the gene’s TSS (±10 kb for promoters; ±100 kb for enhancers) (**Figure S8**; **Table S5**). This maximum distance was chosen to ensure gene regulatory domains were of similar size across species despite disparities in reference genome assembly quality (**Figure S9**). In total, 93%-95% of all one-to-one orthologous genes were associated with at least one active putative CRE in heart, and 95%-96% in liver, depending on the species (**Figure 4B**; 9,073 to 10,989 genes associated with at least one promoter; 12,181 to 12,960 associated with at least one enhancer). This approach assigns a regulatory score to all one-to-one orthologous genes and integrates all CREs within the defined regulatory windows, providing an integrated view of local gene regulatory activity.

**Figure 4:**
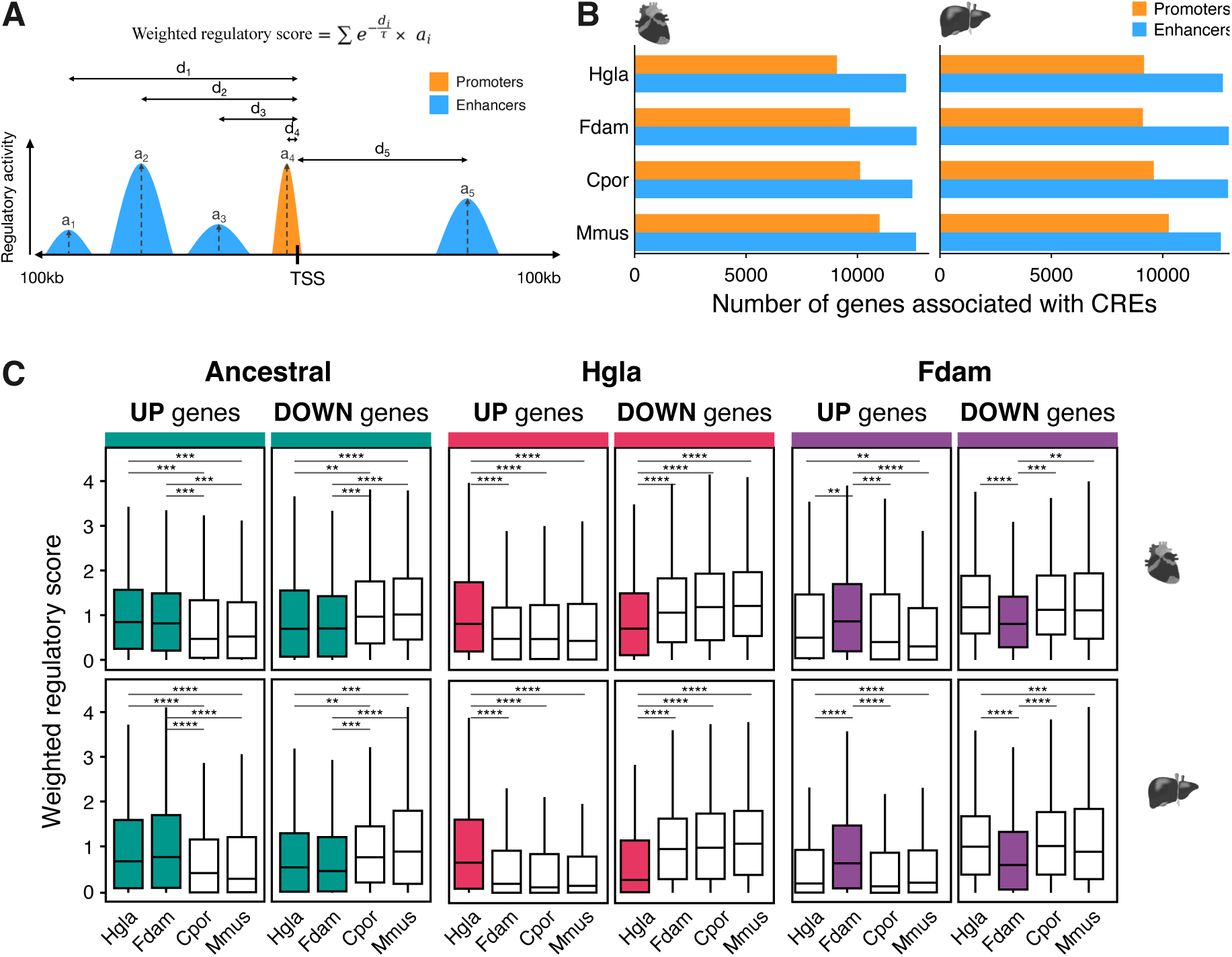
Changes in regulatory activity around TSSs correlate with expression shifts. **(A)** Schematic of the distance-weighted regulatory score calculation. Each cis-regulatory element (CRE) contributes to the gene regulatory score as a function of its activity and distance from the transcription start site (TSS). The contribution of each CRE decreases exponentially with distance from the TSS (**Methods**). **(B)** Numbers of orthologous genes associated with at least one putative promoter or enhancer in heart and liver across species. **(C)** Distributions of normalised weighted regulatory scores for genes with lineage-specific expression shifts. Up-regulated genes show significantly elevated regulatory scores in the foreground branch compared to background species, while down-regulated genes display the inverse pattern.

We first assessed the relationship between gene expression and the regulatory score defined above. As expected from previous studies (Andersson et al. 2014; ENCODE Project Consortium 2012; Kundaje et al. 2015), we found significant positive correlations between expression levels and local CRE activity in all species and tissues (Spearman’s ρ = 0.24–0.38; **Table S6**). To refine this metric, we further implemented a distance-weighted regulatory score in which each CRE’s contribution decays exponentially with its distance to the TSS, with the decay parameter τ chosen to maximise correlation with gene expression in each tissue, similar to previous approaches (Ouyang et al. 2009; Wong et al. 2015; Granja et al. 2021) (**Methods**; **Figure S10A**; **Table S5**). This maximisation improved correlations between regulatory score and expression across species, reaching correlation coefficients of 0.39–0.59 (**Table S5**), indicating that proximal elements contribute most strongly to the correlation with transcriptional output.

We next normalised the dynamic ranges of gene regulatory scores across species (**Methods**; **Figure S10B**; **Table S5**) and examined the distributions of regulatory scores for genes with branch-specific expression shifts. Genes up-regulated in a given species or branch consistently displayed higher regulatory scores in that species compared to others (**Figure 4C**). For instance, up-regulated genes in the naked mole-rat liver showed significantly elevated regulatory activity compared to background species (Wilcoxon’s test, p < 0.0001). Conversely, down-regulated genes showed lower regulatory scores in the foreground species (**Figure 4C**). Notably, no significant differences were observed between background species for either up-or down-regulated genes, with only one exception (up-regulated genes in heart in the Damaraland mole-rat branch, where naked mole-rat showed elevated scores on average compared to mouse). Importantly, these patterns were robust to the regulatory score formula and were also observed with the unweighted regulatory score, which does not maximise correlation with gene expression (**Figure S8C**).

These results indicate genes with species-or branch-specific expression shifts, as detected by our analysis strategy, also exhibit congruent changes in cis-regulatory activity. Moreover, this integrated framework allows us to identify genes in which regulatory and expression changes align, highlighting candidates with compelling evolutionary trajectories that may contribute to adaptive traits.

### Coordinated evidence for lineage-specific accelerated evolution of gene regulation and expression in mole-rats

To further dissect how gene expression evolution may be controlled by regulatory changes, we tested whether genes with expression shifts are more frequently associated with individual CREs that also have shifted activity. In our previous study (Parey et al. 2023), we identified a set of orthologous CREs between all four species with evidence of activity shifts in mole rats, using phylogenetic modelling of ChIP-seq coverage intensity. We defined CREs as “up-shifted” (respectively “down-shifted”) when the phylogenetic model identified a statistically significant increase in H3K27ac signal intensity (respectively decrease) between species consistent with accelerated evolution at that locus. Incorporating this data allowed us to associate orthologous CREs with orthologous genes and examine whether lineage-specific changes in regulatory activity coincide with shifts in gene expression.

We used a closest-gene approach to assign CREs to putative target genes and excluded CRE–gene pairs with very large variations in relative distances between species (greater than ten-fold; **Methods**). This consensus annotation ensured that assignments were comparable across species. We found that 86% of one-to-one orthologous genes were linked to at least one orthologous CRE in liver (70% to at least one promoter and 84% to at least one enhancer) and 87% in the heart (67% to at least one promoter and 81% to at least one enhancer). Across tissues and evolutionary branches, we observed a strong concordance between gene expression shifts and CRE activity shifts. Up-regulated genes were preferentially associated with up-shifted CREs, with odds ratios ranging from 1.1 to 4.1 and highly significant enrichments in nearly all cases (Fisher’s exact test, FDR-adjusted *p* ≤ 4.10^-3^ for all tests except heart enhancers in the ancestral mole-rat branch, which were non-significant; **Figure 5A–B**). Promoters generally showed stronger enrichments than enhancers, particularly in the liver where odds ratios reached 4.1 for promoters and 2.4 for enhancers (naked mole-rat branch: FDR-adjusted *p* = 2.10^-23^ and 1.10^-41^, respectively). Conversely, down-regulated genes were also significantly enriched for associations with down-shifted CREs (odds ratios 1.4–3.4; all FDR-adjusted *p* ≤ 2.10^-2^), with the exception of Damaraland mole-rat promoters in heart (FDR-adjusted *p* = 0.18).

**Figure 5:**
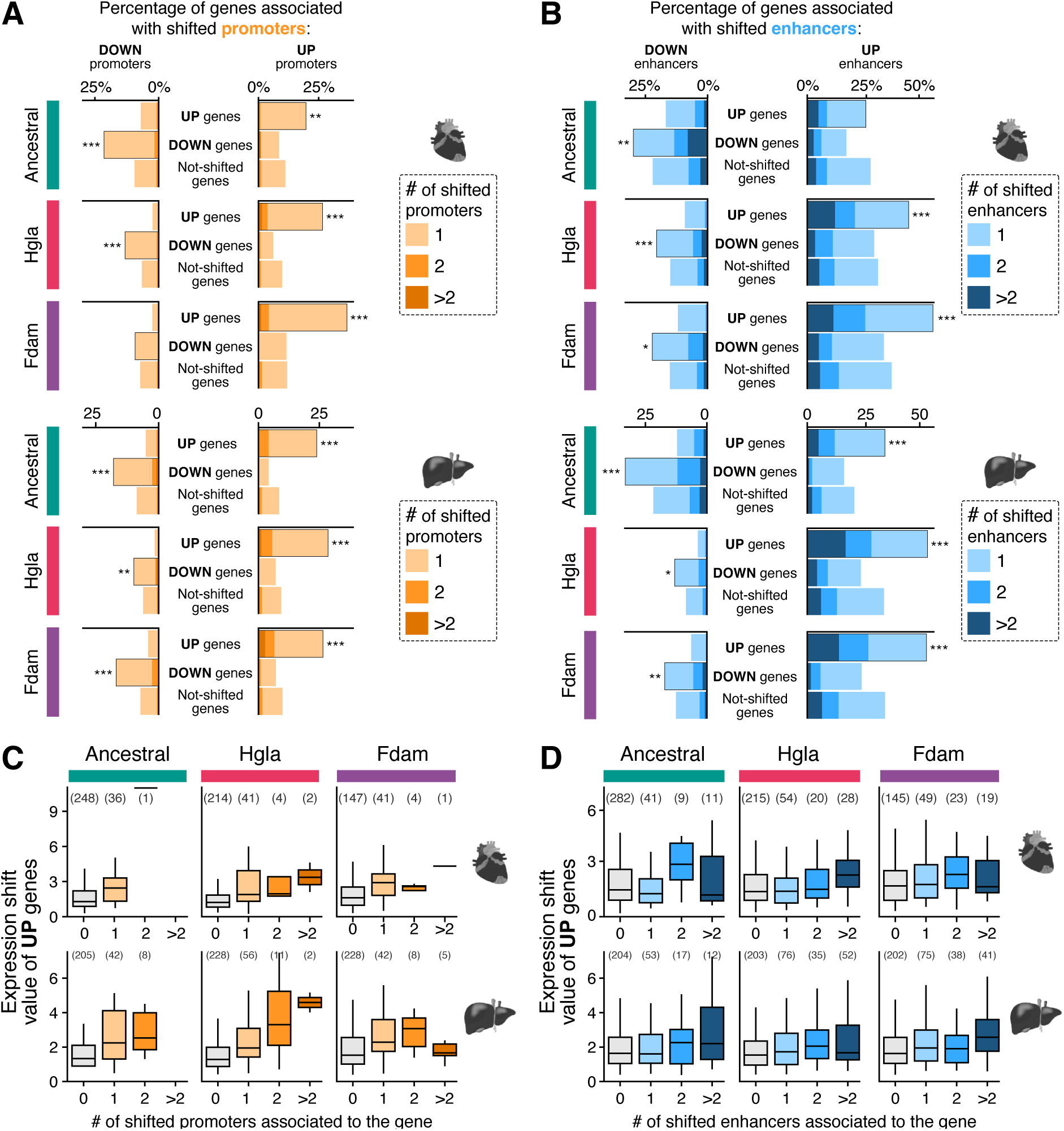
**Concordance between CRE activity shifts and gene expression shifts in mole-rats. (A–B)**. Proportion of shifted genes associated with shifted promoters **(A)** or enhancers **(B)** in heart and liver across evolutionary branches (Ancestral, naked mole-rat, Damaraland mole-rat). Bars show the fraction of genes associated to 0, 1, 2, or >2 shifted CREs, stratified by gene expression status (UP, DOWN, Not-shifted). Genes with expression shifts are significantly enriched for CREs with activity shifts in the same direction. Enrichment was generally stronger for promoters than for enhancers. Asterisks denote statistically significant differences between the highlighted gene group and the other genes (Fisher’s exact test, FDR-adjusted; *: p < 0.05; **: p < 0.01; ***: p < 0.001). **(C–D)** Magnitude of expression shifts in up-regulated genes according to the number of associated shifted promoters **(C)** or enhancers **(D)**. Boxplots show that expression shift values increase with the number of shifted promoters, with significant differences between genes with and without shifted promoters (Wilcoxon test, FDR-adjusted; p < 0.001). Enhancers showed weaker and more variable effects: significant associations were detected only in liver for naked mole-rat and Damaraland mole-rat (FDR-adjusted p < 0.05), whereas no differences were observed in heart.

We next evaluated whether the magnitude of expression divergence scales with the number of up-shifted CREs. We observed positive correlations in all tissues and branches, indicating that genes associated with more shifted CREs tend to exhibit stronger up-regulation (**Figure 5C-D**). The effect was consistently stronger for promoters (e.g. Spearman’s ρ = 0.32 in Damaraland mole-rat heart, Spearman’s ρ = 0.29 in naked mole-rat liver) than for enhancers, where correlations were weaker (e.g. Spearman’s ρ = 0.11 in naked mole-rat heart; Spearman’s ρ = 0.08 in Damaraland mole-rat liver). Consistently, genes with at least one shifted promoter displayed significantly higher expression shift values than those without shifted promoters (Wilcoxon’s test, FDR-adjusted p < 0.001 in all cases). In contrast, enhancer effects were weaker and more variable: significant associations were detected only in liver of naked mole-rat and Damaraland mole-rat (FDR-adjusted p < 0.05), whereas no significant differences were found in heart.

We performed the same analysis for down-regulated genes. Here, no correlation was observed between the number of down-shifted CREs and the magnitude of expression shifts (**Figure S11**), suggesting that cumulative effects are less apparent for negative shifts. Nevertheless, Wilcoxon tests revealed significant associations in several contexts: genes associated with at least one down-shifted promoter or enhancer were significantly down-regulated in the ancestral branch in heart, as well as promoters in heart of naked mole-rat, and in liver in the ancestral and Damaraland mole-rat branches (all FDR-adjusted p < 0.05). These associations were less consistent than for up-regulated genes, suggesting that evolutionary down-regulation of gene expression may rely on different regulatory principles compared to up-regulation.

Together, these findings provide evidence that lineage-specific regulatory changes, captured at the level of single promoters and enhancers, are associated with and quantitatively influence expression divergence. Rather than isolated events, these changes reflect cumulative regulatory dynamics that underpin gene expression evolution. The consistency of this pattern across branches and tissues suggests a generalisable model whereby regulatory element shifts contribute to species-and tissue-specific gene expression changes.

### Concordant transcriptomic and regulatory shifts highlight candidate loci for mole-rat phenotypic evolution

By integrating phylogenetic modelling of gene expression with regulatory activity, our study highlights specific genes with concordant evolutionary trajectories consistent with accelerated evolution (**Table S7**), which may associate with mole-rat adaptations. Here, we describe example loci of particular interest, either because they have been previously associated with phenotypic evolution in mole rats or because they harbour pronounced regulatory shifts and are prime candidates for functional investigation.

First, we highlight genes involved in glycogen metabolism. A previous study reported glycogen metabolism retains neotenic features in naked mole-rats, with the adult heart favouring the liver isozyme of glycogen phosphorylase *Pygl* instead of the muscle isozyme *Pygm* (which typically predominates in mammalian cardiac tissue (Bundgaard et al. 2024). We found that phylogenetic modelling detects this isozyme shift in glycogen phosphorylases and disambiguates how this trait evolved in the mole-rat clade. Indeed, our results suggest *Pygl* was up-regulated in the ancestral mole-rat branch, with the recruitment of several new active enhancers across the gene body that are largely shared between both mole-rat species (**Figure 6A**). Conversely, the muscle isozyme *Pygm* is down-regulated specifically in the naked mole-rat branch, associated with reduced regulatory activity in the local landscape (**Figure 6B**). Together, our results indicate isozyme shifts of glycogen phosphorylase were likely driven by selection and suggest the reduced reliance on *Pygm* in the heart is a further specialization specific to naked mole-rats, while the continued expression of *Pygl* in the adult heart may be shared across mole-rat species.

**Figure 6:**
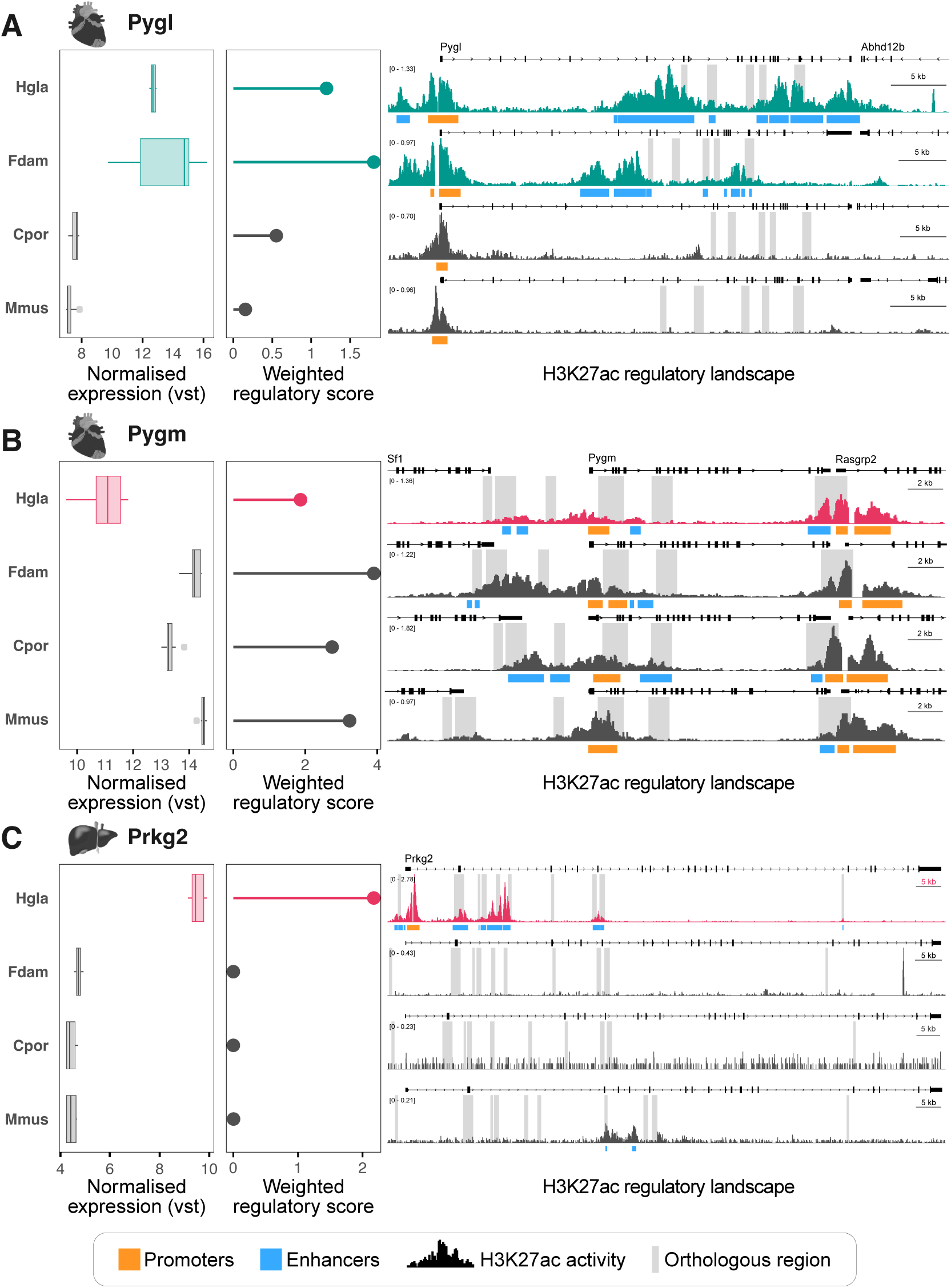
Gene expression, normalised weighted regulatory scores and H3K27ac regulatory landscapes at example candidate loci with concordant transcriptomic and regulatory shifts. (A) Pygl, up-regulated in the ancestral mole-rat branch. (B) Pygm, down-regulated in the naked mole-rat branch. (C) Prkg2, up-regulated in the naked mole-rat liver. Promoters are highlighted in orange and enhancers in blue. One-to-one orthologous regions across the four species are highlited by grey bars, the order of orthologous regions is conserved within each locus.

In the liver, we also identify *Prkg2* as another candidate gene of potential interest with concordant changes in expression and regulation. *Prkg2*, a gene encoding a cGMP-dependent protein kinase involved in metabolic and stress signalling (Flores-Costa et al. 2020), is up-regulated specifically in the naked mole-rat branch and was amongst the top ten genes exhibiting the largest expression shifts in our dataset (**Table S7**). This expression pattern is mirrored by an increased weighted regulatory score and the detection of nine up-regulated enhancers potentially evolving under selection in this branch (**Figure 6C**). Altogether, these threads of evidence suggest accelerated evolution has reshaped gene regulation and expression at this locus during naked mole-rat evolution, and highlights *Prkg2* as a candidate target for further functional studies.

These examples illustrate how the combined analysis of gene expression, regulatory activity, and CRE shifts can uncover lineage-specific changes that align with known physiological traits, while also pointing to additional candidate genes for functional investigation (**Figure S12**).

## Discussion

Gene expression divergence across species is a major contributor to mammalian phenotypic diversity (King and Wilson 1975; Wray 2007; Carroll 2008). This divergence is effected primarily through mutations in non-coding regions that modify the activity of CREs and consequently alter the expression of functionally conserved proteins (Villar et al. 2015; Reilly et al. 2015; Cotney et al. 2013; Roller et al. 2021; Phan et al. 2025). While this relationship is well documented, identifying instances of divergence that may reflect selection rather than drift has been less studied. In this study, we applied a phylogenetic framework to identify lineage-specific shifts in gene expression in African mole-rats and investigate how those changes are supported by their regulatory landscapes. Our primary goal was to identify genes whose expression levels have evolved under adaptive selection rather than diverging due to neutral drift. Detecting selective events on gene expression remains an open challenge in comparative transcriptomics (Price et al. 2022; Bedford and Hartl 2009). Compared to more classical differential expression analyses across species, which conflate drift with adaptive changes, phylogenetic modelling is more discriminative in identifying genes with putatively functional expression differences between species. Further, genes that exhibit both changes in their regulatory landscapes and a significant change in optimal expression levels represent strong candidates for adaptation: our strategy therefore represents a practical roadmap to prioritise genes for functional investigation.

We identified a total of 2,219 and 2,283 unique genes with putative adaptive expression changes in heart and liver, respectively, with an average of 750 per branch and tissue. In contrast, differential expression analysis identified 10,878 unique genes as significantly different between species in at least one tissue (78% of all tested genes), with an average of 6,856 per branch and tissue. This result highlights how phylogenetic modelling strongly improves specificity in detecting putatively adaptive changes in expression levels between species, as well as narrowing down when those changes have likely happened in the evolutionary tree. These findings also suggest that many more genes have experienced putative selective shifts in gene expression compared to positive selection on amino-acid changes at the sequence level: indeed, genes with sequences evolving under positive selection in mole rats range from 142 to 513 genes overall, according to different studies (Sahm et al. 2018; Davies et al. 2015). This aligns with the paradigm that gene regulation likely contributes a substantial fraction of the functional changes underlying species phenotypic divergence, compared to amino-acid substitutions (King and Wilson 1975; Wray 2007; Carroll 2008).

To investigate how regulatory evolution underpins expression divergence, we combined two complementary approaches. The first integrates promoter and enhancer activity into a quantitative regulatory score within a short genomic window around the TSS (about ±20 kb), providing an aggregated measure of local regulatory input. The second considers the activity of orthologous CREs across the four species at short-and mid-range distances from the TSS (up to ±100 kb) and applies phylogenetic modelling to detect lineage-specific activity shifts in these CREs. Those two approaches are complementary, with one capturing the effects of cumulative, quantitative regulatory inputs on transcription, and the other pinpointing individual CREs with evidence of evolutionary changes. Both approaches support that lineage-specific divergence at the transcriptomic level is accompanied by concordant regulatory changes. Earlier reports have shown that conserved expression is typically supported by conserved regulatory activity (Berthelot et al. 2018; Villar et al. 2015; Phan et al. 2025; Wong et al. 2020; Reilly et al. 2015; Sarropoulos et al. 2019): here, we extend this paradigm to show that putatively adaptive shifts in gene expression have a coordinated evolutionary signature that can be jointly detected at the transcriptomic and regulatory levels. Importantly, the correlation we observe between the number of shifted CREs and the magnitude of expression divergence points to cumulative effects of regulatory change, particularly at promoters, on shaping lineage-specific expression profiles. These results emphasise that regulatory and transcriptomic evolutionary dynamics can be harnessed to identify candidate genes that may be involved in phenotypic adaptation.

At the functional level, shifted genes are enriched for biological processes previously implicated in mole-rat physiology, including oxidative phosphorylation (Heinze et al. 2018; Yap et al. 2022), lipid metabolism (Heinze et al. 2018; Farhat et al. 2020), and stress responses (Kim et al. 2011; Xiao et al. 2017). In some cases, mole-rat expression changes in specific genes had been proposed in previous studies, such as for downregulation of respiratory chain components in naked mole-rat liver (Heinze et al. 2018). This analysis strategy relied on detecting coordinated expression shifts across a substantial number of genes in each pathway, and contrasts with previous reports of discrete evolutionary changes in mole-rats associated to phenotypic differences (Kim et al. 2011; Fang et al. 2014; Omerbasic et al. 2016). We therefore combined this approach with prioritisation of candidate loci harbouring signatures of accelerated evolution on both gene expression and regulatory landscape. In particular, we confirm an isoform switch in glycogen phosphorylases: up-regulation of the liver isoform (*Pygl*) in the ancestral mole-rat branch, accompanied by new enhancer activity, and down-regulation of the muscle isoform (*Pygm*) specifically in naked mole-rats. This trajectory supports earlier metabolomic observations of liver-like glycogen use in naked mole-rat heart (Bundgaard et al. 2024) and is compatible with adaptive modification of cardiac metabolism under hypoxic stress. Another example is the up-regulation of *Prkg2* in naked mole-rat liver, mirrored by multiple enhancer gains, which marks it as a promising candidate for further functional investigation. This aligns with previously observed differential expression in the mole-rat liver (Heinze et al. 2018) potentially related to the involvement of Prkg2 acting as a cell proliferation suppressor in the regulation of cellular pathways involved in cancer (Flores-Costa et al. 2020). We also observed accelerated evolution on gene expression and CRE activity for loci involved in oxidative phosphorylation, mitochondrial metabolism (*Ppard*, *Rxra*) or cardiac contraction (*Cacnb1*). In sum, our functional genomics data is consistent with strong selective pressures on gene expression levels and CRE activity in mole-rats – particularly in the heart, to down-regulate processes such as mitochondrial metabolism and cellular respiration.

Our conclusions are constrained by several limitations. First, the phylogenetic sampling in this study is restricted to two mole-rat species and two outgroups, limiting the resolution of evolutionary timing and precluding the detection of complex evolutionary scenarios across the clade. Second, although phylogenetic models aim to detect lineage-specific shifts indicative of positive selection (Rohlfs and Nielsen 2015; Price et al. 2022; Zwonitzer et al. 2023), distinguishing these from relaxation of constraint or compensatory changes in functional genomics data remains an open challenge. Third, functional annotations of candidate genes remain indicative and are largely based on processes described in other clades (in particular mice), which may not be conserved in mole-rats: while our analyses highlight plausible biological processes and candidate loci for adaptation, experimental validation will be required to establish causality and physiological relevance of the described gene expression changes.

Despite these limitations, our study establishes a general framework for dissecting molecular evolution processes in model and non-model species. Extending this approach to broader phylogenies and additional molecular layers, such as 3D chromatin architecture or single-cell resolution datasets, could further refine our ability to detect natural selection acting on gene regulation. Ultimately, by generating phylogenetically-aware hypotheses, this study sets the stage for targeted functional investigations in mole-rat models and beyond, linking regulatory and expression evolution to candidate adaptive traits.

## Methods

### Experimental set up

RNA-seq was used to profile gene expression in heart and liver tissues from four species: the naked mole-rat (*Heterocephalus glaber*; Hgla), Damaraland mole-rat (*Fukomys damarensis*; Fdam), guinea pig (*Cavia porcellus*; Cpor), and mouse (*Mus musculus*; Mmus). Additional RNA-seq replicates were incorporated from previously published datasets: Berthelot et al. 2018 (Berthelot et al. 2018), Bens et al. 2018 (Bens et al. 2018), Faulkes et al. 2024 (Faulkes et al. 2024); with the species and number of replicates detailed in **Table S8**. Across tissues, the number of biological replicates per species ranged from three to eight.

### RNA extraction

Total RNA was extracted from 10-50 mg of tissue samples, previously flash-frozen in dry ice and kept at-80°C. Briefly, frozen tissue samples were homogenised in lysis buffer (RNAEasy, Qiagen) using homogenisation beads (Precellys hard tissue homogenising tubes CK28) in a FastPrep-24 tissue homogeniser (MP Biosciences) with three 20 s pulses at a speed of 4.5 m/s. Homogenised tissue lysates were used directly for RNA extraction using a commercial RNA extraction kit (RNEasy, Qiagen).

### Library preparation for RNA-seq

500 ng of total RNA from each sample were used for preparation of total RNA-sequencing libraries with NEBNext UltraTM II RNA Library Prep Kit for Illumina (New England Biolabs). Briefly, total RNA was depleted of ribosomal RNA (rRNA) with rRNA Depletion Kit v2 (New England Biolabs) and RNA sample purification beads. rRNA-depleted RNA was used for preparation of cDNA and RNA-sequencing libraries following manufacturer specifications, with 12-15 minutes of RNA fragmentation at 94C and 10 cycles of PCR amplification. RNA-sequencing libraries were quantified by quantitative PCR with Kapa Illumina library quantification kit (Roche diagnostics) in a Roche LC480 instrument. RNA-seq libraries were then pooled in equimolar concentrations and sequenced in an Illumina NovaSeq instrument by commercial supplier Novogene UK.

### Genome resources

The reference genome and transcriptomes for naked mole-rat (HetGla1.0), Damaraland mole-rat (DMRv1.0), guinea pig (Cavpor3.0) and mouse (GRCm38) were downloaded from Ensembl v102 (Yates et al. 2020).

### Reannotation of orthologous genes

To increase the number of one-to-one orthologous genes included in the study, we supplemented Ensembl 102 orthology annotation (Yates et al. 2020) with the Zoonomia annotation (Christmas et al. 2023) (**Figure S1**). Zoonomia has released a *de novo* annotation for both mole-rat species and the guinea pig. However, they used different genome assembly references for this *de novo* annotation than those used in Ensembl 102, making direct comparison between Ensembl and Zoonomia’s annotations is impossible. To link Zoonomia’s gene and orthology annotation to Ensembl’s, we conducted a reciprocal BLAST (Altschul et al. 1997) between the transcripts of the two databases.

For each species (naked mole-rat, Damaraland mole-rat and guinea pig), we extracted the FASTA sequences of all Ensembl 102 annotated transcripts using the `getfasta` function of the bedtools suite (Quinlan and Hall 2010) and used them to produce a BLAST database. We replicated this process on the annotated one-to-one orthologous transcripts from Zoonomia.

Next, we performed a reciprocal BLAST, where Ensembl transcripts were blasted against the Zoonomia database and vice versa. Note that Zoonomia used the following assemblies for their de novo gene annotation: cavPor3 from Ensembl 80 for the guinea pig, DMR1.0_HiC from UCSC for the Damaraland mole-rat, and HLhetGla3 *de novo* assembled by Zoonomia for the naked mole-rat.

For each BLAST, we extracted the best hit. Transcript pairs that matched in both BLASTs were retained. The overall percentage of identity of the BLASTs hit retained was at 99.9% with a minimum of 76%. This method allowed us to identify 2,226 additional one-to-one orthologous genes. The final set of one-to-one orthologous (14,408 genes) comprised of the 12,182 Ensembl ono-to-one orthologous genes and 2,226 ono-to-one orthologous genes from Zoonomia.

### RNA-seq processing

#### Quality control and trimming

We performed a quality control of the RNA-seq libraries using FastQC (www.bioinformatics.bab raham.ac.uk/projects/fastqc). All the libraries passed this quality control. Subsequently, we removed sequencing adaptors and low-quality reads using Cutadapt (Martin 2011) with the following parameters: a minimum read length of 25 bp after trimming (`-m 25`), a minimum sequence quality score of 30 (`-q 30`), and the removal of any reads containing more than one N base (`--max-n 1`).

#### Expression quantification and normalisation

We next focused on quantifying gene expression and applying normalisation. This step is critical in cross-species transcriptomic comparisons to mitigate technical biases, balance discrepancies in sequencing depth, and account for biological variability. Without proper normalisation, spurious signals of selection may be detected. A key source of biological variability to consider is the difference in lengths of orthologous genes and transcripts across species, which can confound comparisons of expression profiles. To address this, we implemented two complementary strategies.

In strategy 1, we normalised raw counts for library size and gene length, following a standard approach in comparative genomics (Brawand et al. 2011; Berthelot et al. 2018; Cardoso-Moreira et al. 2019). In strategy 2, we used transcript-level quantification from Salmon, which corrects for effective transcript length rather than gene length. This approach more precisely accounts for length differences and incorporates transcript diversity, thereby mitigating discrepancies in annotation quality among species. For both strategies, we generated principal component analyses (**Figure S3**), which revealed that strategy 2 improves replicate clustering within species and tissues and better captures the phylogenetic signal.

**Strategy 1:** Gene length normalisation

**STAR alignment and featureCounts gene quantification:** Trimmed reads from each species were aligned to the corresponding reference genome (see *Genome resources*) using STAR v2.7.10 (Dobin et al. 2013). We then quantified gene-level counts with featureCounts v2.0.1 (Liao et al. 2014) against the Ensembl v102 annotation.

**Counts normalisation:** For each species, we imported raw counts of one-to-one orthologous genes into DESeq2 v1.36 (Love et al. 2014). We extracted gene length information from the Ensembl v102 annotation and incorporated it into the DESeq2 object using the `avgTxLength` parameter. We then normalised counts for both gene length and library size using the DESeq function. The combined dataset across species was normalised with the median-of-ratios method, which adjusts for sequencing depth by estimating sample-specific size factors as the median ratio of observed counts to a pseudo-reference (the geometric mean across samples for each gene). To control for technical variation introduced by different data sources, we included a batch term in the DESeq2 design formula (∼species + data source). Finally, we applied a variance-stabilising transformation (VST) to the normalised counts for downstream visualisation.

**Strategy 2:** Effective transcript length normalisation

**Salmon alignment and transcript quantification:** Trimmed reads from each species were pseudo-aligned and quantified to their respective reference transcriptome (see *Genome resources*) using Salmon v.013.1 (Patro et al. 2017), with a decoy-aware index. The pseudo-mapping process included corrections for GC content bias (`--gcBias`) and sequence-specific biases (`--seqBias`). Quantification was performed at the transcript level.

**Counts normalisation:** For each species, we summarised transcript-level abundance estimates generated by Salmon to the gene level using the tximport v1.24 R package (Soneson et al. 2016), with counts corrected for effective transcript length using the lengthScaledTPM method. To enable cross-species comparisons, we retained only one-to-one orthologous genes (see “*Reannotation of orthologous gene*”). We then merged gene-level count matrices across species and imported them into DESeq2 (Love et al. 2014). We normalised the combined dataset across all samples using the median-of-ratios method implemented in DESeq2, which adjusts for library size by computing sample-specific size factors as the median of ratios of observed counts to a pseudo-reference (the geometric mean across samples for each gene. Finally, variance-stabilising transformation (VST) was applied to the normalised counts for downstream visualisation.

Downstream analyses were performed using normalised counts from strategy 2.

### Differential gene expression analysis

We performed a differential expression analysis on the normalised counts using DESeq2.). To account for technical variation introduced by different data sources, we included a batch term in the DESeq2 model to account for data sources (∼species + data source). For each tissue, we tested three contrasts corresponding to the foreground lineages of interest: (i) naked and Damaraland mole-rats versus mouse and guinea pig (ancestral mole-rat branch), (ii) naked mole-rat versus the other three species (naked mole-rat branch), and (iii) Damaraland mole-rat versus the other three species (Damaraland mole-rat branch). Genes were considered differentially expressed if their Benjamini–Hochberg adjusted p-value was below 0.05. Results of the differential analysis are reported in **Table S3**.

### Phylogenetic modelling of expression shifts with EVE

We used the normalised counts obtained from strategy 2 (log2-transformed with a pseudocount of 1; **Table S2**) as input to model the evolution of gene expression in three specific branches using the EVE model (Rohlfs and Nielsen 2015). Specifically, we applied the `twoThetaTest` function from the evemodel R package (Gilad et al. 2006) to test for expression shifts in the ancestral mole-rat branch, the naked mole-rat branch, and the Damaraland mole-rat branch. To compute empirical p-values, we performed 1,000,000 simulations under the null hypothesis, following the recommendations of Rohlfs and Nielson 2015 (Rohlfs and Nielsen 2015). Simulations were carried out using code adapted from Parey et al. 2023 (Parey et al. 2023). We used this high number of simulations to ensure precision in empirical p-value estimation, as the number of unique values scales with the number of simulations. Finally, for each branch and tissue, we intersected the list of significant genes identified by EVE (empirical p-value< 0.05) with those obtained from the differential analysis performed with DESeq2 (see above) and retained only genes present in both lists. Results of EVE analysis are reported in **Table S3**.

### Functional enrichment analysis

We performed a Gene Set Enrichment Analysis (GSEA) using the fgsea v1.22.0 R package (Bioconductor, n.d.). Gene sets were downloaded from the Mouse Molecular Signatures Database (Liberzon et al. 2015) using the msigdbr v7.5.1 R package, including collections from GO Biological Processes (GO BP) and Reactome. For each branch and tissue, we performed a GSEA analysis, ranking genes based on EVE empirical p-value signed by shift value. Enrichment scores were calculated using the `fgsea` function, with a minimum gene set size of 10 genes and a maximum gene set size of 500 genes. The fgsea package also conducts a leading-edge analysis to identify the core subset of genes contributing to the enrichment signal. Gene sets were considered significantly enriched if their BH-adjusted p-values were below 0.05. Results of this analysis are reported in **Table S4**.

### Cis-regulatory element activity quantification

For each species and tissue, H3K27ac ChIP-seq and matched input BAM files were downloaded from Parey et al. 2023 (Parey et al. 2023). Regulatory element activity was quantified with featureCounts v2.0.1 (Liao et al. 2014), using the sets of promoters and enhancers identified by Parey et al. 2023. Read counts were converted into RPKM values separately for H3K27ac and input libraries. Regulatory activity was then estimated as the ratio of H3K27ac over input RPKM, with input values floored at 1 to avoid inflation at low coverage. The resulting activity values were quantile-normalised across replicates and species and log₂-transformed within each tissue and species.

### Unweighted and weighted regulatory score calculation

To summarise the regulatory activity of genes, we computed a window-based regulatory score for each gene by summing the activity of cis-regulatory elements (CREs) within fixed windows around the transcription start site (TSS). Promoters were defined as active CREs located within ±10 kb of the TSS, while enhancers were defined as active CREs located within ±100 kb of the TSS (**Figure S8A**). These maximal distances were chosen to ensure coverage of alternative promoters and proximal regulatory elements, which can extend beyond the immediate TSS region in many genes. For each gene, we assigned all overlapping CREs based on these distance thresholds and calculated the regulatory score as the sum of their H3K27ac activity values (see previous method section).

To refine the regulatory score, we implemented a distance-weighted metric that sums the contributions of all associated promoters and enhancers while weighting them according to their genomic distance from the transcription start site (TSS). Each CRE contributes to the score according to the formula ∑𝑒 *e*^−^*^d^*τ ×*a* where d is the distance of the CRE to the gene’s TSS, a is the regulatory activity of the CRE, and τ is a decay parameter controlling the effect of distance. To identify an optimal τ value, we tested a grid of τ values ranging from 1 to 60,000. For each species and tissue, we computed the correlation between gene expression levels and the corresponding weighted regulatory scores (**Figure S10A**). For each case, the τ yielding the maximum correlation was retained, with correlation values rounded to two decimals to account for plateaus where several τ values performed equivalently. To ensure comparability across species within the same tissue, we then selected the largest τ value among those giving maximal correlations across species. This approach avoids bias towards species-specific optima while maximising the cross-species robustness of the metric. Following this procedure, the chosen τ values were 4000 for the heart and 3750 for the liver. To enable comparisons across species and tissues, raw regulatory scores were quantile-normalised within each species and tissue prior to downstream analyses (**Figure S8B** and **S10B**).

Unweighted and weighted regulatory scores are reported in **Table S5**.

### Orthologous CREs assignment to genes

Orthologous CREs and their shift direction were obtained from Parey et al. 2023 (Parey et al. 2023). To assign orthologous CREs to target genes, we used a distance-based approach consistent with the strategy applied for the regulatory score calculation. Promoters were assigned to genes within ±10 kb of the transcription start site (TSS), and enhancers within ±100 kb. The 100 kb window for enhancers was chosen to ensure comparable regulatory domains across species given the differences in genome assembly quality among non-model species.

Because the analysis was restricted to orthologous CREs, we excluded gene–CRE pairs for which the relative distance was not conserved across species. To quantify this, we calculated a conservation score based on the number of species in which the CRE–gene distance did not deviate by more than 10-fold from the maximum allowed distance (100 kb for promoters, 1 Mb for enhancers). Gene–CRE pairs with a conservation score below three (i.e. in which more than one species had the CRE located further than 10-fold the maximum distance from the gene) were excluded. Out of the initial 233,487 pairs, 222,000 passed this filter, with 90% of pairs conserved across all four species. In total, this annotation included 27,948 promoter–gene pairs and 81,948 enhancer–gene pairs in heart, and 30,802 promoter–gene pairs and 81,514 enhancer–gene pairs in liver. This resulted in a consensus region-to-gene annotation independent of any single species, providing a good framework for cross-species comparisons. CRE-gene pairs are reported in **Table S9**.

### Regulatory track visualisation

Genome browser tracks were generated from H3K27ac ChIP-seq BAM files for each species and tissue. Coverage was computed using bedtools genomecov (Quinlan and Hall 2010), with read counts scaled to one million mapped reads to allow comparisons across samples. For each species and tissue, replicate coverage tracks were averaged and converted into bigWig format for visualisation.

## Acknowledgements

We thank Leo Zeitler and all the members from the Berthelot Lab for discussion and feedback. We thank David Gaynor and Katy Goddard (Kalahari Research Trust), and Nigel Bennet (University of Pretoria) for technical assistance; Tim Clutton-Brock (Kalahari Research Trust) for the permission to collect tissues from Damaraland mole-rats; and Ewan St. John Smith (University of Cambridge) and Thomas Park (University of Illinois at Chicago) for provision of naked mole-rat samples. Finally, we thank Jane Reznick for reading the manuscript and providing valuable feedback.

## Author contributions

CB, DV and MD designed the study. DV performed experiments.

MD performed computational analyses.

EL and EP provided key feedback and discussions.

MD, CB and DV wrote the manuscript with input from EL and EP.

## Data availability

RNA-sequencing data reported here has been deposited to the Gene Expression Omnibus repository with accession number GSE304970. We also included in the study previously published datasets (accession numbers GSE222972 and EMTAB-4550). Pipelines and scripts used to perform the analyses are available at https://gitlab.com/maelle.daunesse/daunesse_et_al_2025.

