## Supplementary Figures for "Coordinated shifts in gene expression and regulation during mole-rat evolution"

#### Table of supplementary figures

### Supplementary figures

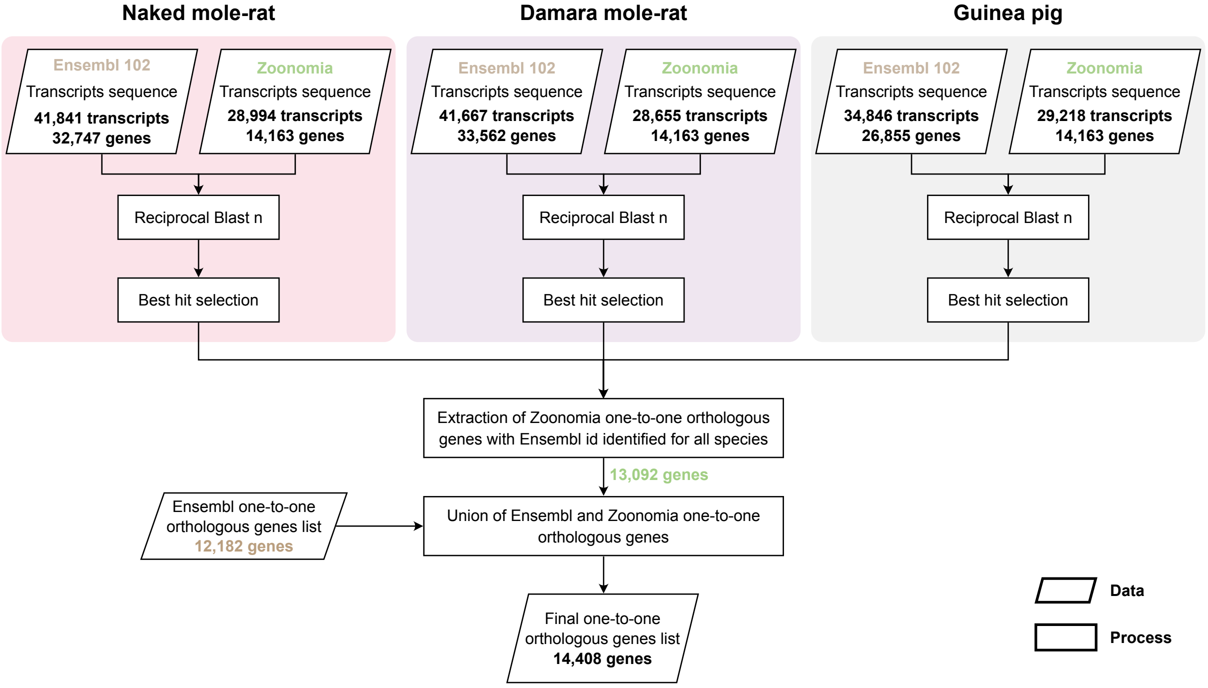

***Figure S1: Orthology reannotation.***

*Workflow of orthology reannotation combining Ensembl v102 and Zoonomia datasets. Reciprocal BLASTs between transcript sets recovered additional one-to-one orthologues, resulting in 14,408 genes retained across the four species.*

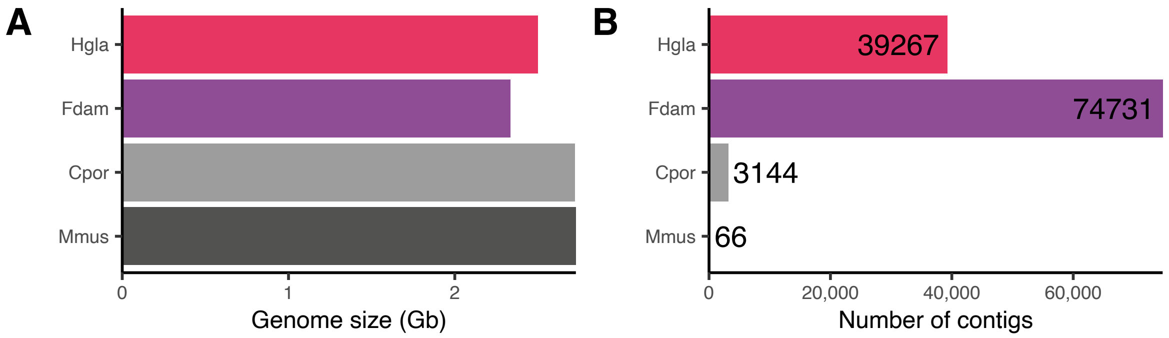

***Figure S2: Genome assembly quality.***

*Comparison of **(A)** genome size and **(B)** number of contigs per genome across the four species (Hgla, Fdam, Cpor, Mmus)*

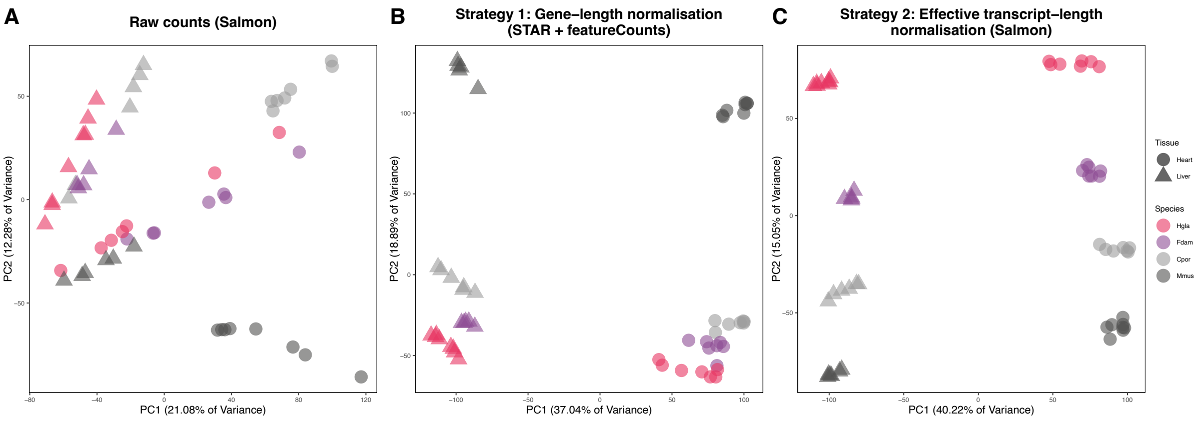

***Figure S3: Normalisation strategy comparison.***

*Principal component analysis (PCA) of expression matrices using (A) raw counts and counts normalised by (B) theoretical gene length (strategy 1) or (C) effective transcript length (strategy 2). Effective-length normalisation (strategy 2) improves replicate clustering and enhances the phylogenetic signal.*

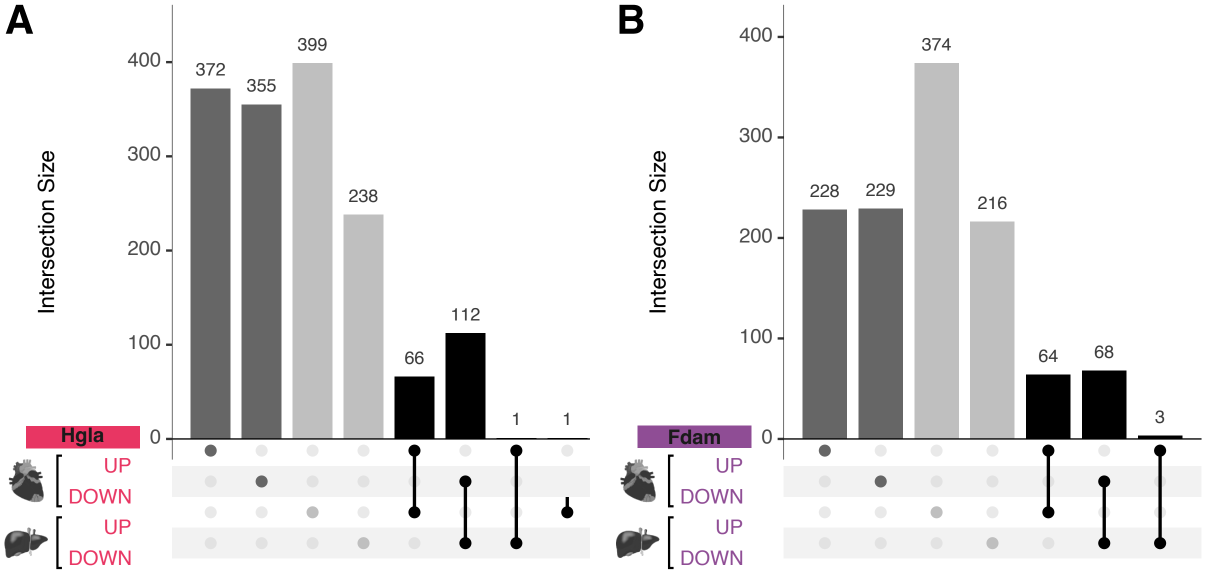

***Figure S4: Overlap of shifted genes between tissues.***

*Upset plot showing the intersection of genes with expression shifts detected by the EVE model in liver and heart for the naked mole-rat (Hgla) and Damaraland (Fdam) branches. Shared genes represent tissue-shared lineage-specific shifted genes.*

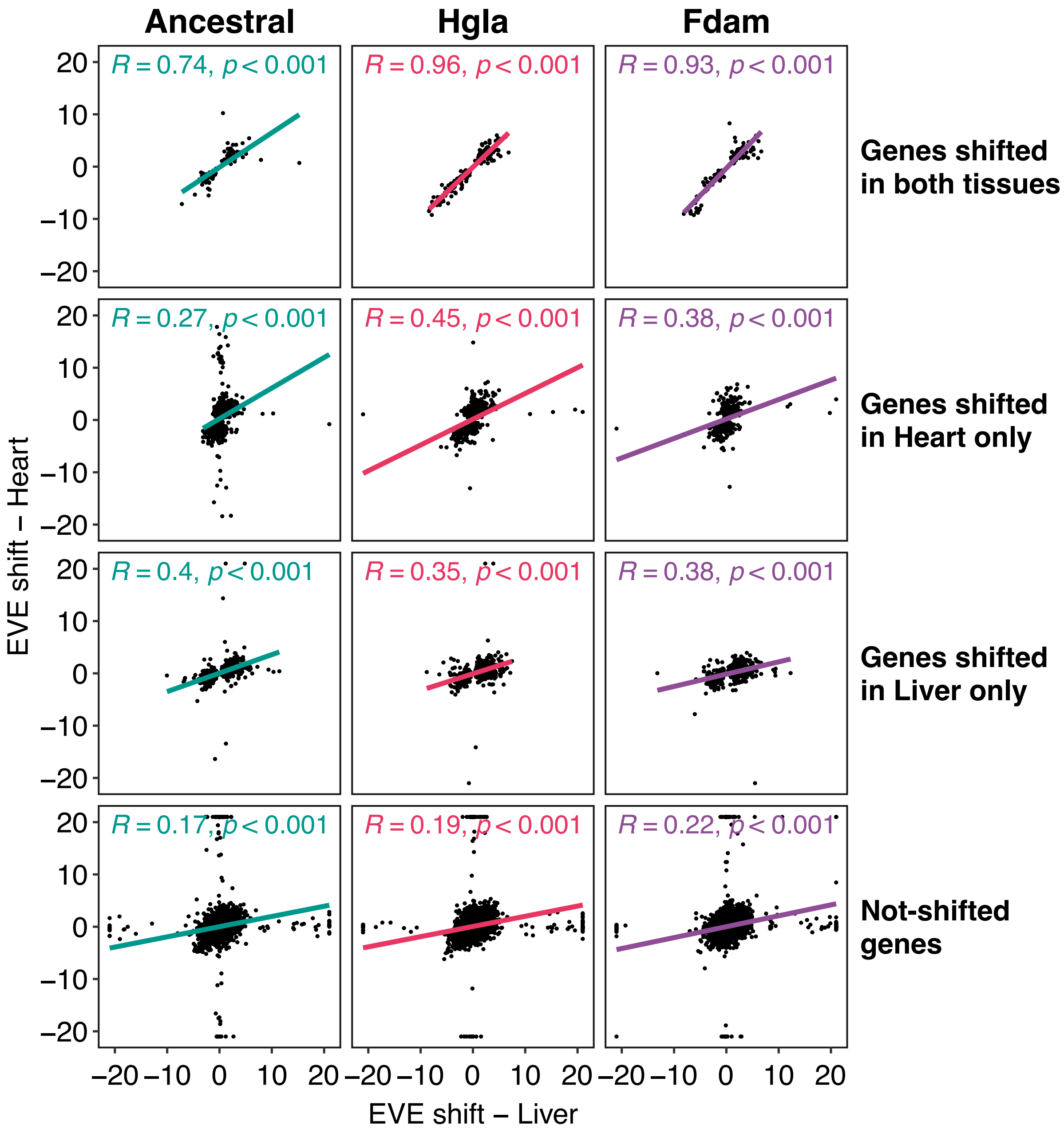

***Figure S5: Correlation of shift magnitudes.***

*Scatterplots showing correlation of EVE shift magnitudes for genes shifted in both tissues (top row), in Heart only (2nd row), in Liver only (3rd row) and in neither tissues (last row). High correlation coefficients indicate consistency of expression shift amplitude across tissues.*

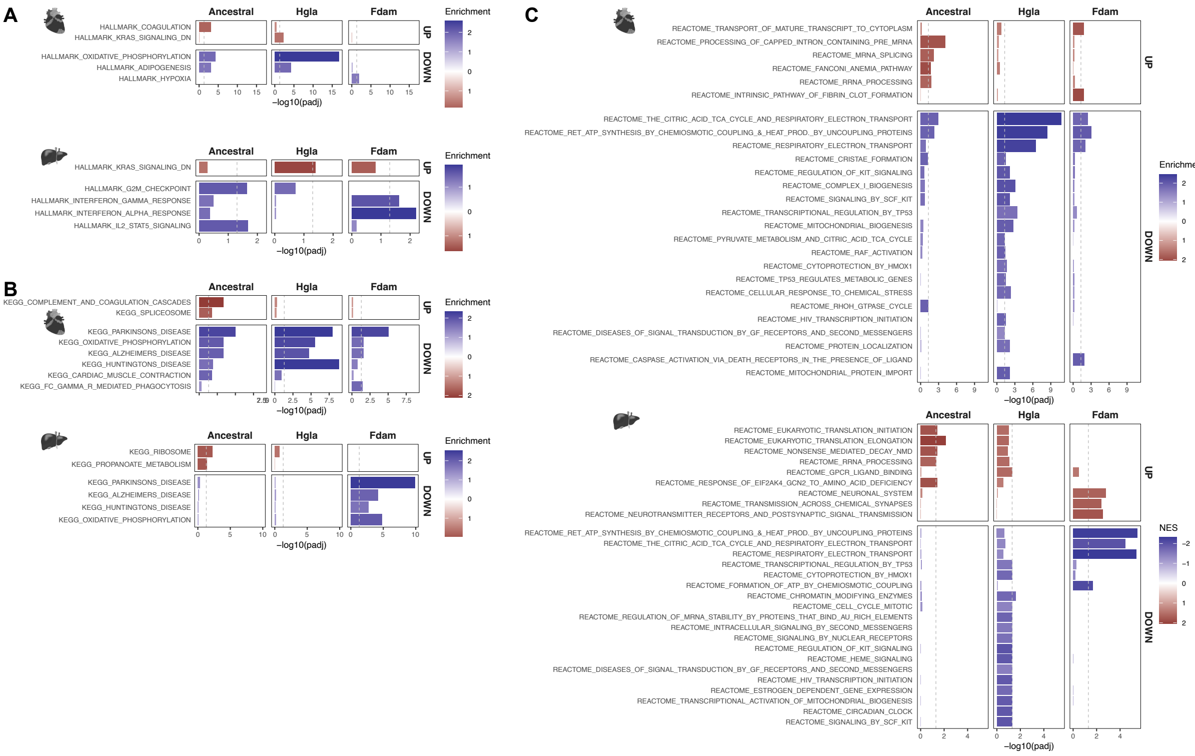

#### *Figure S6: Functional enrichment of shifted genes.*

*Gene set enrichment analysis (GSEA) of shifted genes for each branch and tissue, using **(A)** Hallmark, **(B)** KEGG and **(C)** Reactome collections. The significantly enriched terms for up-regulated (UP) and down-regulated (DOWN) genes are shown, coloured by normalised enrichment score (red: enriched in up-regulated genes; blue: enriched in down-regulated genes).*

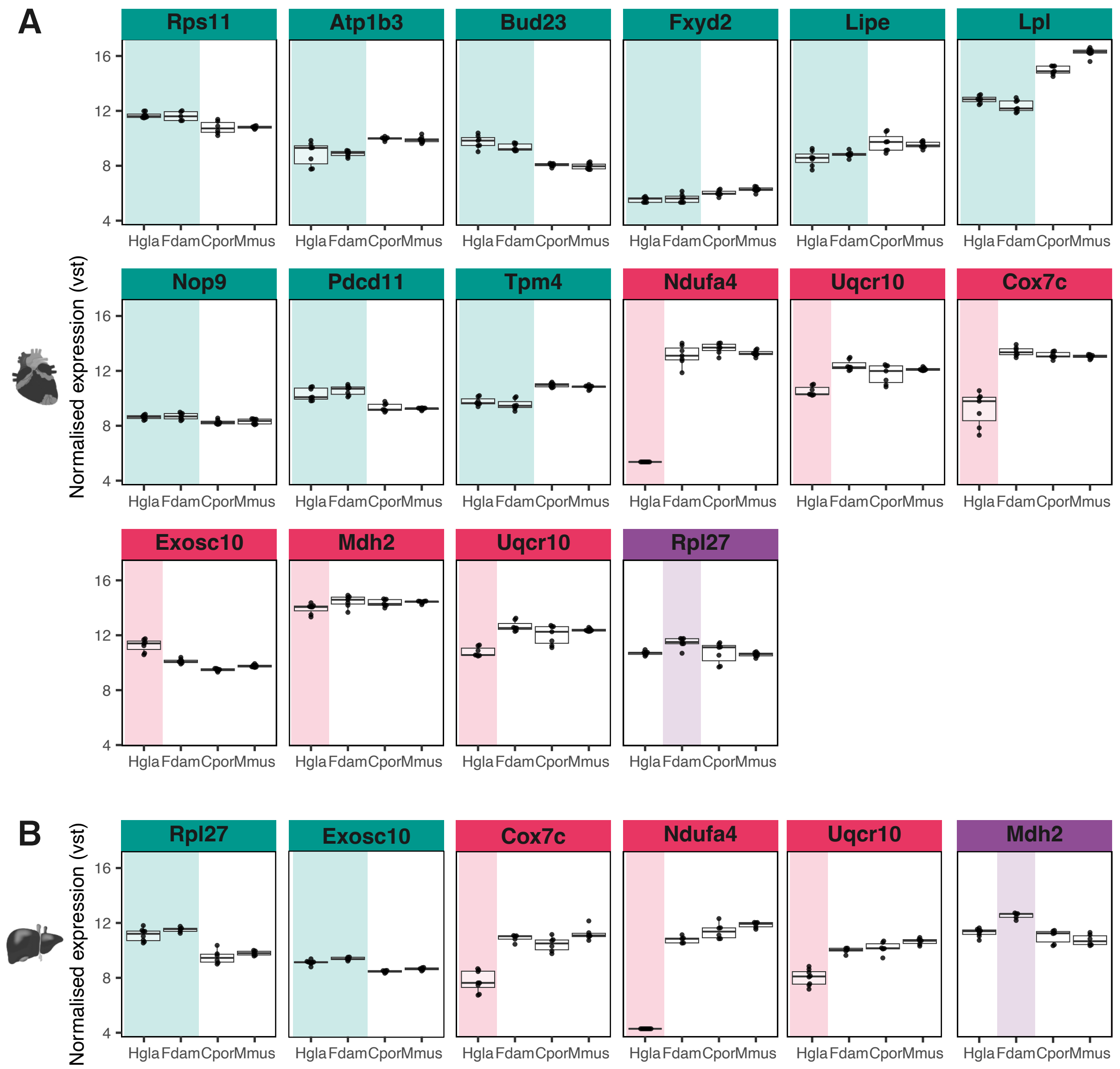

***Figure S7: Example expression profiles of genes associated with significantly up- or down-regulated gene sets.***

*Variance-stabilised expression levels of genes belonging to gene sets significantly enriched in the GSEA analysis (see corresponding GO terms in **Figure 3C** and **Figure S6**). Boxes represent interquartile ranges with median lines. Genes in turquoise are shifted in the ancestral mole-rat branch, genes in pink are shifted in the naked mole-rat branch and genes in purple are shifted in the Damaraland mole-rat branch.*

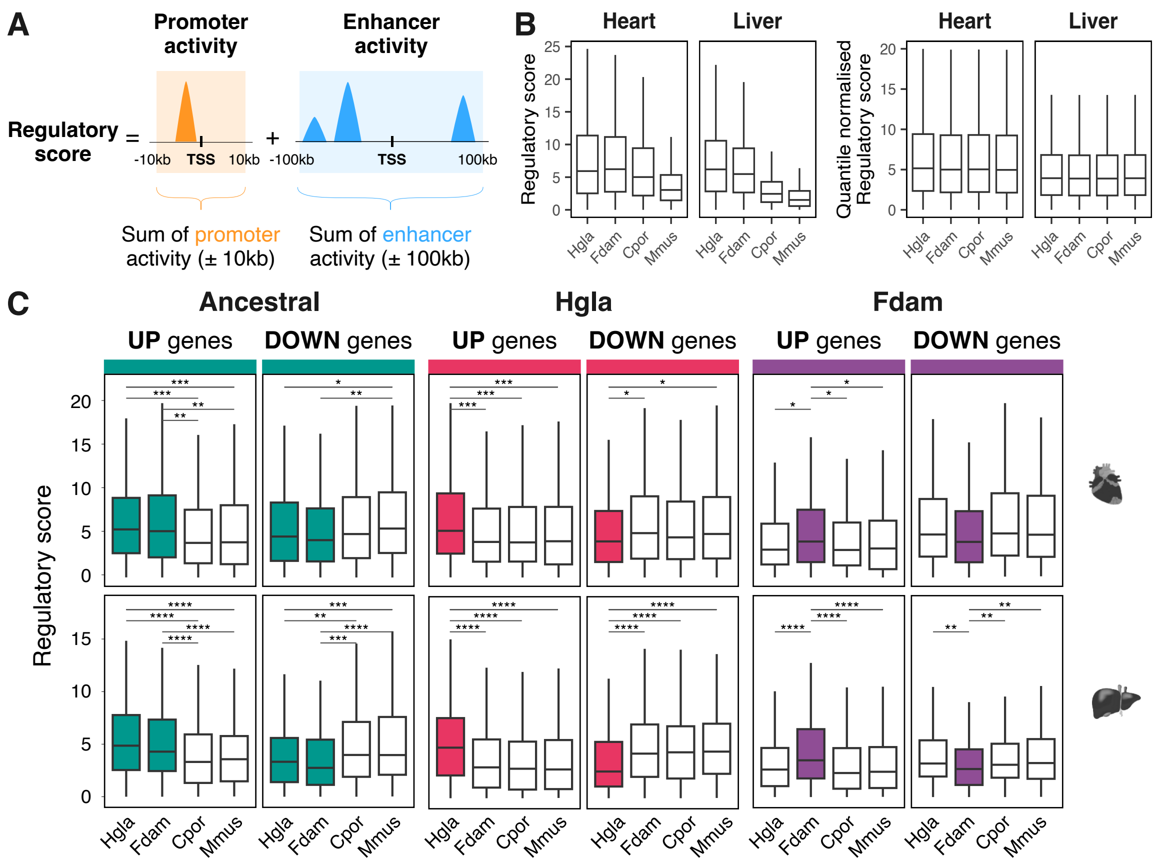

***Figure S8: Relationship between regulatory scores and expression shifts.***

***(A)*** *Schematic representation of regulatory score calculation corresponding to the sum of promoter activity (orange; ±10 kb from the TSS) and enhancer activity (blue; ±100 kb from the TSS). Each cis-regulatory element (CRE) contributes to the total score according to its H3K27ac signal intensity within the specified window.* ***(B)*** *Regulatory score distribution before and after quantile normalisation.* ***(C)*** *Distributions of normalised regulatory scores for genes with lineage-specific expression shifts. Up-regulated genes show significantly elevated regulatory scores in the foreground branch compared to background species, while down-regulated genes display the inverse pattern.*

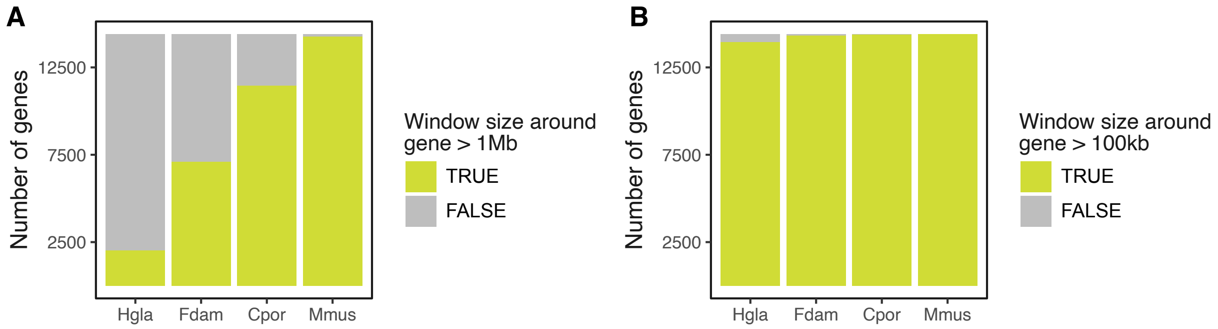

***Figure S9: Window size for regulatory element assignment to gene.***

***(A)*** *Number of gene with a ±10 Mb around TSSs.* ***(B)*** *Number of gene with a ±100 kb around TSSs.*

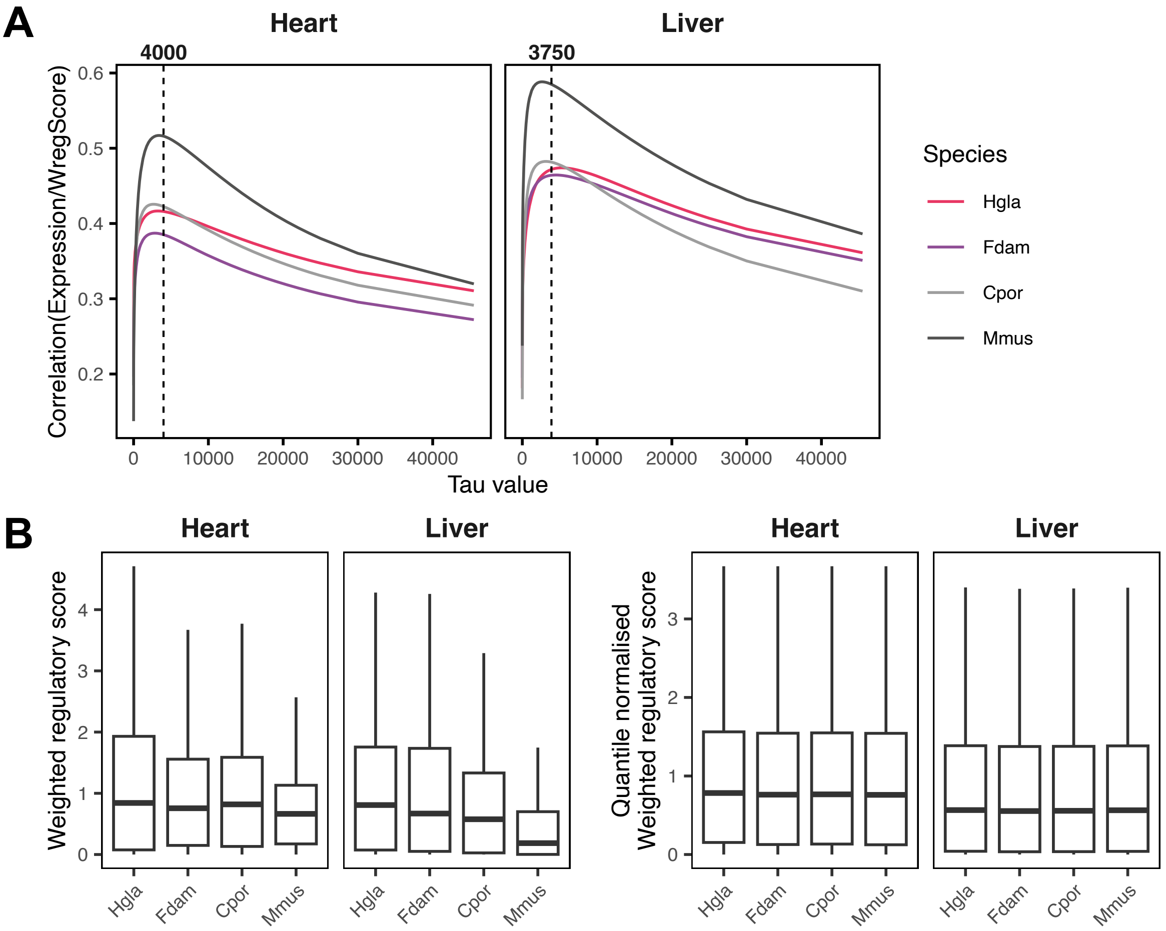

***Figure S10: Optimisation of the weighted regulatory score.***

***(A)*** *Correlation between gene expression and regulatory activity as a function of distance from the transcription start site (TSS). Each line represents the Spearman correlation coefficient (ρ) between expression and weighted regulatory score for a range of distance-weighting parameter (τ) going from 0 to 40,000. Dashed line represent the selected tau for each tissue (4000 for Heart and 3750 for Liver).* ***(B)*** *Weighted regulatory score distribution before and after quantile normalisation.*

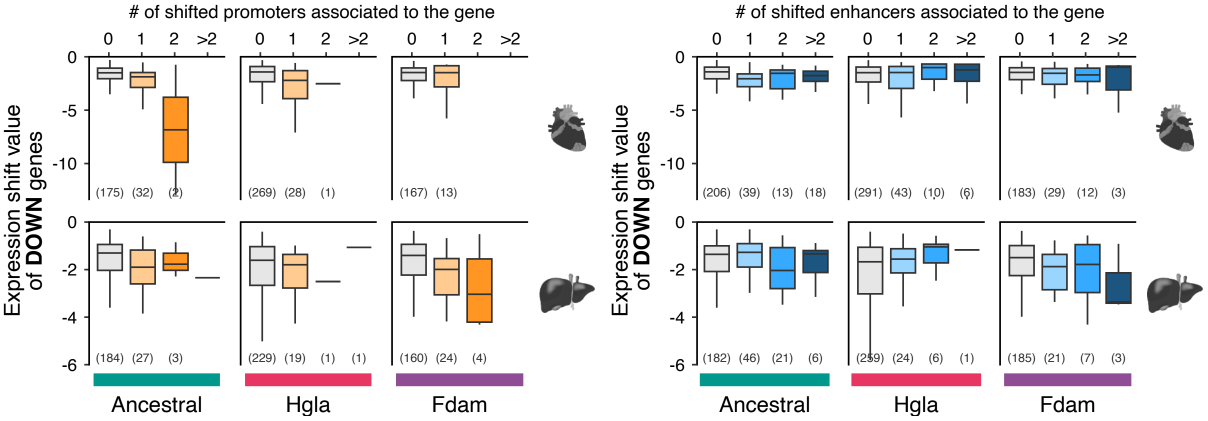

*__Figure S11: Magnitude of expression shifts in down-regulated genes.__*

*Magnitude of expression shifts in down-regulated genes according to the number of associated shifted promoters (__left__) or enhancers (__right__).*

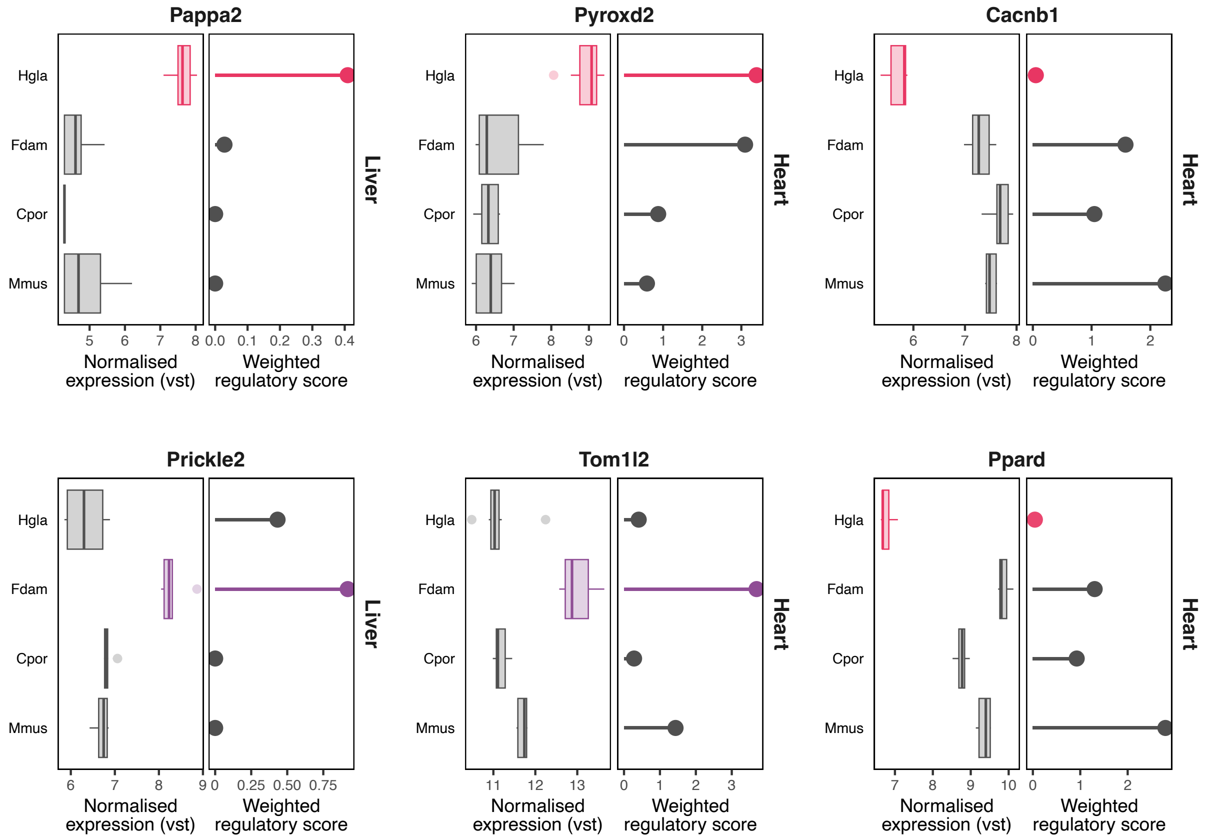

*__Figure S12: Gene expression, normalised weighted regulatory scores at example candidate loci with concordant transcriptomic and regulatory shifts.__*

### Supplementary tables

***Table S1: List of 1-to-1 orthologous genes across the four species.***

*(see excel file named: S1_ortholog_gene_list.xlsx).*

***Table S2: Heart and Liver normalised expression counts.***

*(see excel file named: S2_normalised_counts.xlsx).*

***Table S3: Results of differential expression (DESeq2) and phylogenetic modelling (EVE) analyses, and their intersection.***

*(see excel file named: S3_supp_shifted_genes.xlsx).*

***Table S4: Results of Gene Set Enrichment Analysis (GSEA) performed with fgsea****.*

*(see excel file named: S4_GSEA_results.xlsx).*

***Table S5: Regulatory score and weighted-regulatory score values per gene****.*

*(see excel file named: S5_regulatory_score.xlsx).*

| Tissue | Species | cor(expression/regScore) | cor(expression/WregScore) |
|---|---|---|---|
| Heart | Hgla | 0.28 | 0.42 |
| | Fdam | 0.24 | 0.39 |
| | Cpor | 0.26 | 0.42 |
| | Mmus | 0.3 | 0.52 |
| Liver | Hgla | 0.34 | 0.47 |
| | Fdam | 0.34 | 0.46 |
| | Cpor | 0.34 | 0.48 |
| | Mmus | 0.38 | 0.59 |

***Table S6: Correlation between regulatory scores and gene expression across tissues and species.***

*Spearman correlation coefficients between gene expression levels and two regulatory score metrics: the window-based regulatory score (regScore) and the distance-weighted regulatory score (WregScore). Correlations were computed separately for each tissue and species.*

***Table S7*****: Gene shift information at the expression and regulatory level**.

(see excel file named: S7_geneCREshift.xlsx).

| | | Replicate # | New data | Berthelot et al. 2018 | Faulkes et al. 2024 | Bens et al. 2018 |
|---|---|---|---|---|---|---|
| **Liver** | **Hgla** | **8** | 4 | 4 | | |
| | **Fdam** | **5** | 5 | | | |
| | **Cpor** | **6** | 3 | 3 | | |
| | **Mmus** | **6** | 2 | 4 | | |
| **Heart** | **Hgla** | **7** | 3 | | 4 | |
| | **Fdam** | **7** | 5 | | 2 | |
| | **Cpor** | **7** | 3 | | | 4 |
| | **Mmus** | **8** | 3 | | 5 | |

***Table S8: Number of replicates per species and tissue with source of the data.***

*Table S9*: **Region to gene output**.

(see excel file named: S9_region2gene.xlsx).
